# Cryo-EM structures of human RNA polymerase III in its unbound and transcribing states

**DOI:** 10.1101/2020.06.29.177642

**Authors:** Mathias Girbig, Agata D. Misiaszek, Matthias K. Vorländer, Aleix Lafita, Helga Grötsch, Florence Baudin, Alex Bateman, Christoph W. Müller

## Abstract

RNA polymerase III (Pol III) synthesises tRNAs and other short, essential RNAs. Human Pol III misregulation is linked to tumour transformation, neurodegenerative and developmental disorders, and increased sensitivity to viral infections. Pol III inhibition increases longevity in different animals but also promotes intracellular bacterial growth owing to its role in the immune system. This highlights the importance to better understand human Pol III transcription on a molecular level. Here, we present cryo-EM structures at 2.8 to 3.3 Å resolution of transcribing and unbound human Pol III purified from human suspension cells that were gene-edited by CRISPR-Cas9. We observe insertion of the TFIIS-like subunit RPC10 into the polymerase funnel, providing insights into how RPC10 triggers transcription termination. Our structures also resolve elements absent from *S. cerevisiae* Pol III such as the winged-helix domains of RPC5 and an iron-sulphur cluster in RPC6, which tethers the heterotrimer subcomplex to the Pol III core. The cancer-associated RPC7α isoform binds the polymerase clamp, potentially interfering with Pol III inhibition by the tumour suppressor MAF1, which may explain why overexpressed RPC7α enhances tumour transformation. Finally, the human Pol III structure allows mapping of disease-related mutations and might contribute to developing inhibitors that selectively target Pol III for therapeutic interventions.

---

Eukaryotes utilise Pol III to transcribe tRNAs, the spliceosomal U6 small nuclear RNA (U6 snRNA) and other short, abundant RNAs^1^. Thus, Pol III must be tightly regulated by oncogenes and tumour suppressors, including the Pol III-specific repressor MAF1^2^. Pol III comprises 17 subunits that form a conserved catalytic core, the stalk domain and Pol III-specific subcomplexes, which are homologous to general transcription factors in the Pol II system^3^ (Fig. 1a). The TFIIE/F-like RPC3-RPC6-RPC7 heterotrimer functions in transcription initiation^4–6^ and the TFIIF-like RPC4-RPC5 heterodimer is required to initiate^7,8^ and terminate transcription^9,10^. Pol III also contains the TFIIS-like subunit RPC10 that is essential for termination and facilitates recycling of Pol III^11,12^, but its precise function in these processes is poorly understood.

**Fig. 1.**
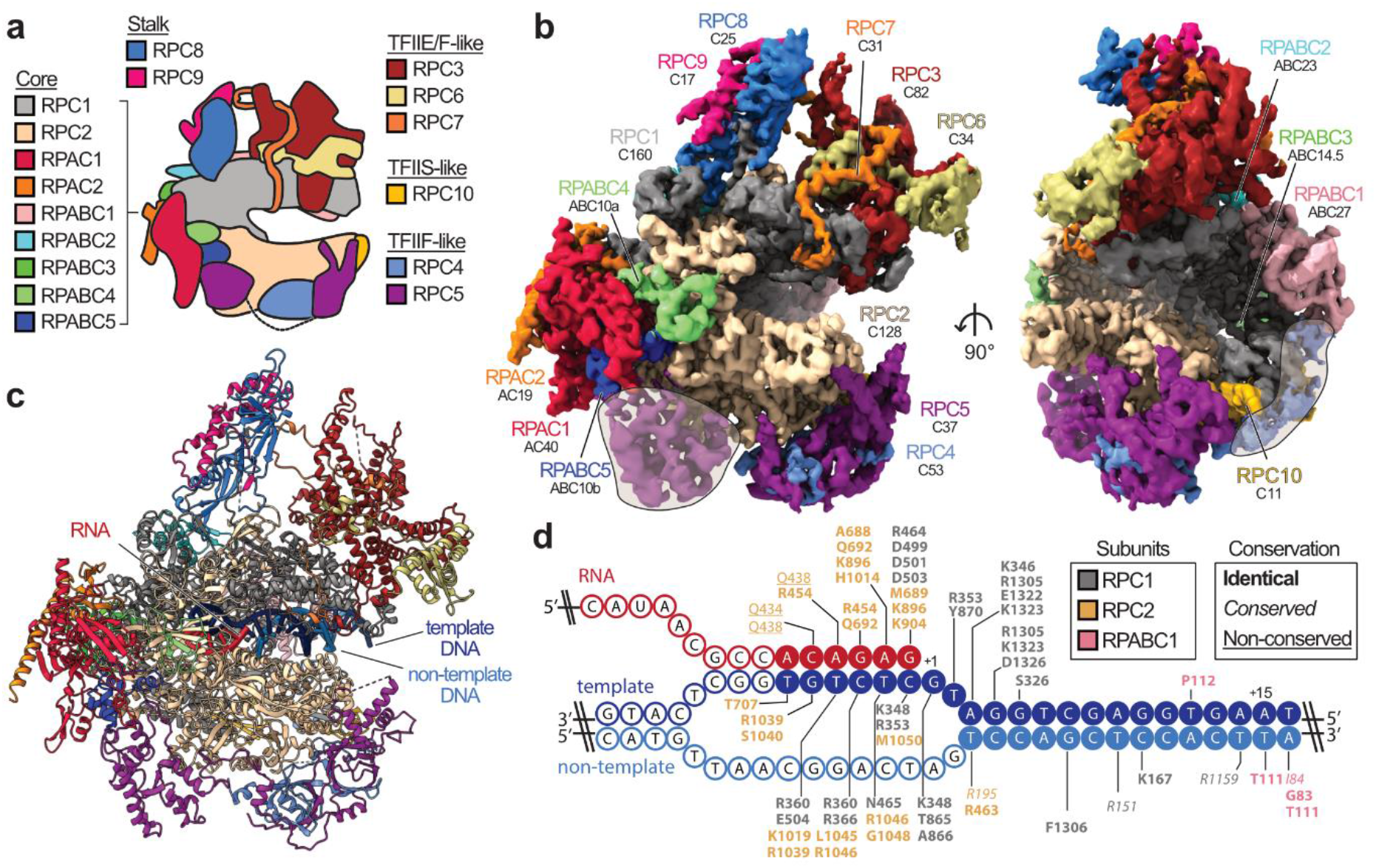
Cryo-EM structure of human RNA polymerase III. **a**, Schematic of human Pol III and colour code of its subunits grouped by subcomplexes or homology to Pol II factors. **b**, Cryo-EM map of apo human Pol III shown from the side (left) and front (right) with human subunits coloured as in **a** and additionally labelled with their yeast counterparts below. Shadings mark superimposed features of a cryo-EM map derived from a pooled dataset of apo and elongating Pol III. **c**, Structural model of elongating human Pol III with bound nucleic acids. **d**, Schematic of the DNA/RNA transcription scaffold. Filled and unfilled circles represent modelled and non-modelled nucleotides, respectively, labelled with contacting residues within 4 Å distance. Boxes on the top-right show the subunit colour-code and used typography of the residues to mark their conservation between human and yeast Pol III.

In humans, the Pol III-specific subcomplexes harbour additional features that are absent in *S. cerevisiae*. RPC5 is twice as large as its yeast counterpart C37^7^, RPC6 comprises an iron-sulphur cluster^13^ of unknown function, and human Pol III can utilise different isoforms of RPC7 (also known as POLR3G and hRPC32)^14^. Whereas RPC7β is ubiquitously expressed, RPC7α is enriched in tumour cells and undifferentiated embryonic stem cells^14^. Ectopic expression of RPC7α also enhances tumour transformation^14^.

Cryo-electron microscopy (cryo-EM) studies on yeast Pol III revealed its architecture^15^ and shed light on the mechanisms of Pol III transcription initiation^16–18^ and Maf1-mediated repression^19^. However, a mechanistic understanding of Pol III function in human health and disease has been hampered due to the lack of an atomic human Pol III structure. Here, we present cryo-EM structures of human Pol III at 2.8 to 3.3 Å resolution in different functional states. Our results expand the overall knowledge of Pol III transcription and provide molecular insights into how higher eukaryote-specific features and disease-causative mutations modulate human Pol III activity.

## RESULTS

### Structure of human Pol III

We used CRISPR-Cas9 to generate a homozygous HEK293 suspension cell line carrying a mCherry-StrepII-His-tag on the C-terminus of subunit RPAC1 (Extended Data Fig. 1a,b) to purify the endogenous 17-subunit Pol III complex. This yielded intact, homogenous and transcriptionally active human Pol III, which we confirmed by negative-stain EM, mass spectrometry and RNA primer extension experiments (Extended Data Fig. 1c-e). We acquired cryo-EM data for Pol III alone (apo Pol III) (Extended Data Fig. 2a,b) and bound to a DNA/RNA scaffold mimicking the transcription bubble, and, thus, resembling the elongating Pol III complex (EC Pol III) (Fig. 1d, Extended Data Fig. 2c). Apo Pol III, was resolved at a nominal resolution of 3.3 Å with the core reaching up to 3 Å (Fig. 1b, Extended Data Figs. 2b, 4a,c). The cryo-EM map of EC Pol III could be refined to 2.8 Å nominal resolution and extended to 2.5 Å local resolution in the core (Extended Data Figs. 2c, 4a,d). More flexible regions could be resolved by focused classification, which yielded, in total, eight partial cryo-EM maps (Extended Data Figs. 3, 4b) that were used to build the atomic models of human Pol III in four functional states: apo Pol III, EC-1 Pol III (RPC5 in conformation 1), EC-2 Pol III (RPC5 in conformation 2), and EC-3 Pol III (RPC10 ‘inside funnel’) (see below). We, hereafter, refer to the EC-1 Pol III structure when describing structural aspects of the elongating human Pol III complex. The structural models contain all 17 subunits forming the core (RPC1, RPC2, RPC10, RPAC1, RPAC2, RPABC1-5), the stalk (RPC8, RPC9), the heterotrimer (RPC3, RPC6, RPC7) and the heterodimer (RPC4, RPC5) (Fig. 1a,c). In elongating human Pol III, the downstream double-stranded DNA (dsDNA) that unwinds into the non-template- and template strand in the active centre is well resolved. The template strand forms a hybrid with the transcribed RNA, of which six bases were built (Fig. 1d, Extended Data Fig. 4h). Nucleic acid-binding is mediated by the core subunits RPC1, RPC2 and RPABC1 (Fig. 1d). Most residues that bind the nucleic acids are identical or conserved between yeast and human Pol III (Fig. 1d), as it has also been observed for mammalian Pol II^20^. Interestingly, the strongest differences between yeast and human Pol III are located in the Pol III-specific heterotrimer and heterodimer subcomplexes, as discussed in more detail below.

### RPC10 swings into the active site

Our cryo-EM maps capture different conformations of the mobile domain of the TFIIS-like subunit RPC10, and of the RPC4-RPC5 heterodimer. Pol III utilises both elements to autonomously terminate transcription on poly-thymidine DNA sequences^9,11^. This termination mechanism is unique to Pol III and critical for its function in the innate immune system by sensing viral DNA^21,22^. RPC10 is anchored to the polymerase core via an N-terminal zinc ribbon domain^15^ (N-ribbon), followed by a flexible linker and a C-terminal zinc ribbon domain (C-ribbon) that is homologues to the Pol II elongation factor TFIIS (Fig. 2a). Similarly to TFIIS, RPC10 uses an acidic hairpin (D88, E89) for hydrolytic 3’ RNA cleavage, and deletion of the acidic hairpin is lethal in *S. pombe*^11^. Our cryo-EM reconstruction of elongating human Pol III reveals that the C-ribbon of RPC10 can adopt two conformations – outside and inside the polymerase funnel (Fig. 2b, Extended Data Figs. 5a,b, 6a,b). In the ‘outside funnel’ conformation, the flexible linker adopts a kink whereby the C-ribbon folds back and contacts the central part of the linker and the RPC1 jaw via its tip (Fig. 2b, Extended Data Fig. 6a,e). The contacted residues are not conserved between yeast and human, explaining why this conformation has not been observed in the yeast Pol III structures^15,16^ (Extended Data Fig. 6c) and suggesting a species-specific adaption to RNA cleavage (Extended Data Fig. 6e). In agreement to this, RNA-cleavage activities of human and yeast Pol III are comparable at 37 °C and 28 °C, respectively (Extended Data Fig. 6f). In contrast, the yeast enzyme cleaves RNA faster than human Pol III at 37 °C. The linker extends when adopting the ‘inside funnel’ conformation (EC-3 Pol III), which is accompanied by a ~110° rotation and a ~57 Å movement of the acidic hairpin (Fig. 2b). Consequently, the C-ribbon inserts into the polymerase pore and funnel, and its position resembles that of the C-ribbon in previous TFIIS-Pol II structures^23–25^ but the acidic hairpin does not reach the 3’ RNA end (Extended Data Fig. 6b,d). To further explore the conformational dynamics of the RPC10 C-ribbon, we subjected our cryo-EM reconstructions to 3D variability analysis (3DVA)^26^. 3DVA of elongating human Pol III reveals that the insertion of the RPC10 C-ribbon into the funnel is accompanied by an ‘open clamp’ conformation (Fig. 2b, Supplementary Video 1). Multiple studies reported that Pol III requires RPC10 (C11 in yeast) to terminate transcription, but that its 3’ RNA cleavage activity is not essential for this process^9,11,12,27,28^. We propose that the insertion of RPC10 C-ribbon induces partial opening of the mobile clamp domain and, thereby, supports the release of transcribed RNA from a paused pre-termination complex previously reported^10^. RPC10-induced clamp opening could also explain why in yeast, this subunit is critical for facilitated re-initiation of transcription^9,29^ because an open clamp has been suggested to be required for binding the closed dsDNA of the promoter^16,17^. In that regard, the release of RPC10 from the active site could potentially also assist promoter DNA melting during the transition from an open to a closed clamp conformation.

**Fig. 2.**
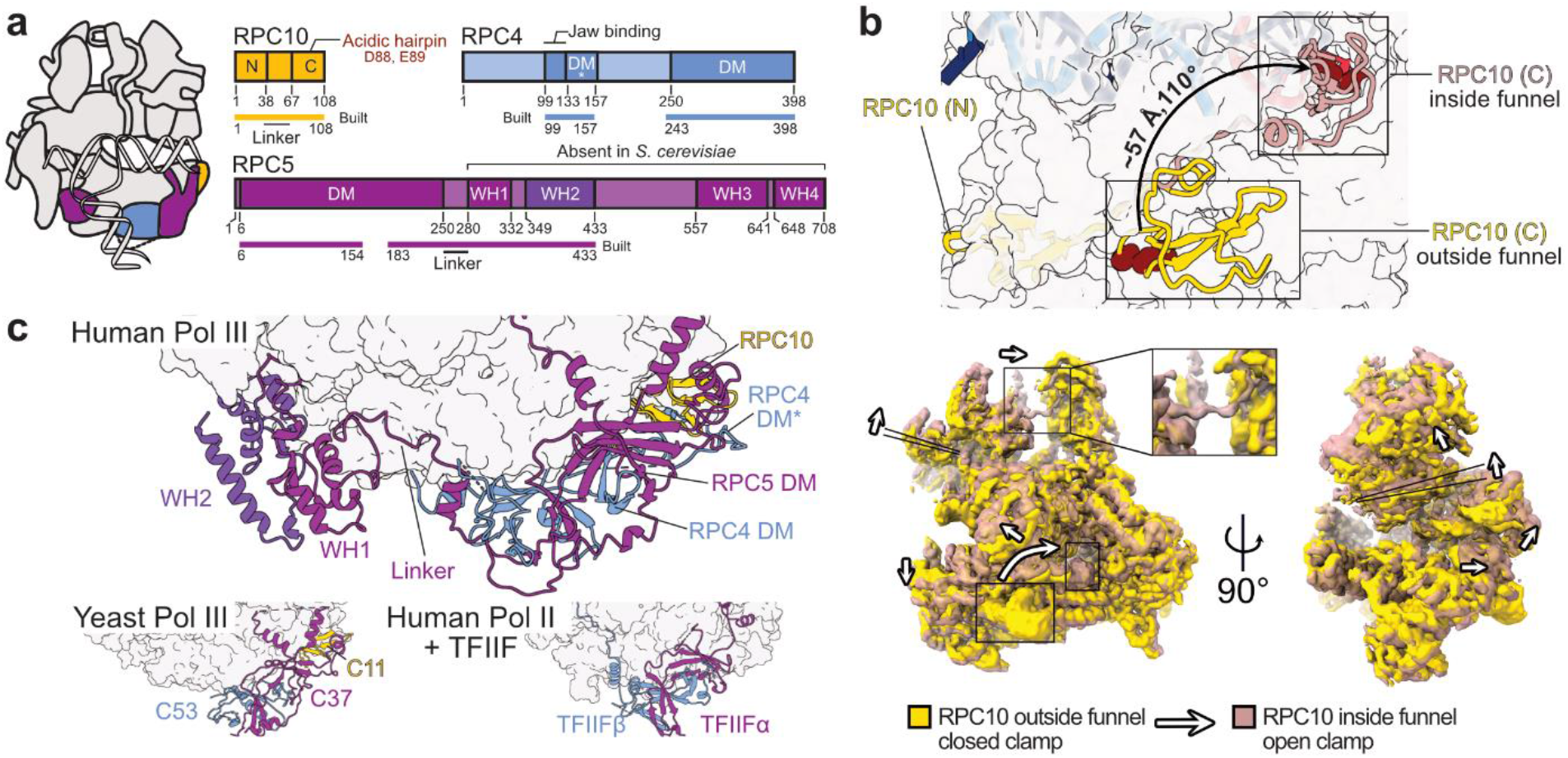
Conformational dynamics and extensions of RPC10 and RPC4-RPC5. **a**, Schematic domain organisation of subunits RPC10, RPC4, and RPC5 in the elongating human Pol III complex. Coloured bars indicate modelled regions and are labelled as ‘built’. N – N-terminal zinc ribbon; C – C-terminal zinc ribbon; DM – dimerization module; WH – winged-helix domain. **b**, Conformational switch of RPC10. Top: Pol III core is shown as transparent surface and RPC10 as cartoon in two conformations of its C-terminal zinc ribbon domain (C): outside (yellow) and inside (light red) the Pol III funnel. The black arrow illustrates the movement of the acidic hairpin (D88, E89), shown as dark red spheres, to engage the active site. Bottom: Elongating human Pol III in two superimposed conformations derived from 3D Variability Analysis (3DVA) with RPC10 outside (yellow map) and inside (light red map). Positions of the C-ribbon are boxed. Conformational rearrangements are highlighted with white arrows and indicate the partial opening of the clamp domain. The framed close-up view shows the appearance of the connection between heterotrimer and stalk in the ‘open clamp’ conformation. **c**, Position of the RPC5 WH1 and WH2 domains, which are connected by a flexible linker to the DM of RPC4-RPC5 in the human Pol III EC (top). RPC4-RPC5 binds polymerase core via its DM domain in a similar fashion as C53-C37 in yeast Pol III (bottom left, PDB 5fj8) and TFIIF in the human Pol II PIC (bottom right, PDB 5iy6). Labelled subunits are shown as cartoons and polymerase cores as surfaces.

### Extended structure of human RPC4-RPC5

RPC4 and RPC5 (C53 and C37 in *S. cerevisiae*) assemble via a dimerization module (DM) and bind the polymerase core in a similar fashion as C53-C37 in yeast Pol III^15^ and TFIIFα/β in the human Pol II pre-initiation complex (PIC)^30^ (Fig. 2a,c). Human RPC4 contains an extended DM (termed DM*, 133-157) that further runs along the RPC1 jaw domain and enters the DNA binding cleft (Extended Data Fig. 5c). Although not visible in our 3D reconstruction, the RPC4 N-terminal part (1-99) is predicted to bind DNA (Extended Data Fig. 7a), raising the possibility that it binds the downstream DNA in the context of another functional state such as the human Pol III PIC. The yeast orthologue C53 is a major target for post-translational modification by the small ubiquitin-like modifier (SUMO) pathway, which represses Pol III activity^31^. Similarly to yeast C53, human RPC4 also has been shown to be sumoylated (Extended Data Fig. 7a). Whereas most of the reported SUMO-modified residues lie in a flexible region (158-249) that is not visible in our structure, we could map one of these residues (K141) onto the newly built DM* region. This lysine is buried inside the RPC4 DM* - RPC5 DM - RPC10 (N-ribbon) interface (Extended Data Fig. 7b) and, thus, can only be targeted for sumoylation if the RPC4-RPC5 dimer unfolds. This could serve as a signal to mark misfolded human Pol III for proteasomal degradation.

RPC5 is the least conserved Pol III subunit, and its size largely varies across species. The C-terminal extension region spanning residues 280-708 is predicted to harbour four winged-helix (WH) domains that are absent in *S. cerevisiae* C37 (Fig. 2a, Extended Data Fig. 7a,e). The cryo-EM density reveals that the WH1 and WH2 domains of RPC5 are closely intertwined and bind the Pol III core between subunits RPC2, RPAC1, and RPABC5 (Fig. 2c, Extended Data Fig. 7c). The two WH domains are connected to the RPC5 DM domain via a flexible linker. As revealed by masked 3D classification, the WH1-2 domains can also adopt a second conformation (EC-2 Pol III) and can point towards the downstream promoter DNA in the modelled human PIC (Extended Data Fig. 7d,f,). The WH3 and WH4 domains are not visible in the cryo-EM reconstructions, indicating that they are mobile and suggesting a function in transcription initiation. Similar functions are reported for the WH domains of yeast Pol I (A49 subunit)^32^, human Pol II (RAP30 subunit of human TFIIF)^30^ and yeast Pol III (C34 subunit)^16–18^. The majority of residues in RPC5 are not predicted to bind DNA (Extended Data Fig. 7a), suggesting that the RPC5 WH domains interact with basal transcription factors instead. The absence of the RPC5 WH domains in *S. cerevisiae* C37 prompted us to ask if these domains co-evolved with a set of basal transcription factors that are also missing in yeast: TFIIIB subunit Brf2 and the SNAP complex (SNAPc), which are both required for U6 snRNA transcription in humans^33^. A comprehensive phylogenomic analysis of RPC5, Brf2 and SNAPc subunits reveals that WH1-4, Brf2 and functional SNAPc are mostly absent in fungi whereas nematodes and flies harbour both WH1-2 and SNAPc but lack WH3-4 and Brf2 (Extended Data Fig. 7e, Supplementary Table 1). In contrast, vertebrates and plants encode all of these genetic elements, possibly suggesting a co-evolution and cooperative action of the RPC5 WH domains with Brf2 and the SNAPc. In agreement to this, modelling of the human Pol III PIC indicates that the WH domains are positioned in a way that would allow interactions with Brf2 and SNAPc (Extended Data Fig. 7f). However, the phylogenomic analysis also shows that this proposed function would not hold true for some species like the parasite *Leishmania major* (Extended Data Fig. 7e) suggesting another or additional function, such as contribution to complex assembly or stability.

### The FeS domain ties heterotrimer and core

The heterotrimer RPC3-RPC6-RPC7 subcomplex (C82-C34-C31 in *S. cerevisiae*) sits on top of the polymerase clamp and functions in transcription initiation, for which it utilises multiple conserved WH domains of RPC3 and RPC6^15^ (Fig. 3a). Our cryo-EM structure shows that the human- and yeast heterotrimer bind the polymerase clamp in a similar fashion (Fig. 3b, Extended Data Fig. 8b). In addition, human RPC6 coordinates a previously described cubane 4Fe-4S cluster^13^ involving four cysteines in its C-terminal region that forms a globular domain (FeS domain) and binds the RPC1 clamp (Fig. 3b,c, Extended Data Fig. 8b,d). The FeS domain is sandwiched between RPC3, RPC7 and the RPC1 clamp and, thereby, serves as an additional interaction hub that interconnects the heterotrimer subunits and ties them to the polymerase core. In agreement with this observation, the 4Fe-4S cluster has been shown to facilitate dimerization and polymerase clamp engagement of the archaeal TFEα/β transcription factor, which is homologous to TFIIE and RPC3-RPC6^13^. Interestingly, the FeS domain is missing in *S. cerevisiae* and most, except four, budding yeast species, but is otherwise widely conserved (Extended Data Fig. 8a). To answer how budding yeasts compensate for the loss of the FeS domain, we compared the human heterotrimer structure with the one from *S. cerevisiae*. The comparison reveals that *S. cerevisiae* RPC3 (C82) features two extensions that bind the polymerase core and are missing in the other analysed model organisms (Extended Data Fig. 8c). The visible part of the N-terminal extension (31-46) overlaps with the position of the RPC6 FeS domain (Extended Data Fig. 8d). The central insertion (331-354) is sandwiched between the clamp and a second helix-turn-helix extension (202-258) of *S. cerevisiae* RPC3 (C82) that binds the downstream DNA and is also absent in the human heterotrimer (Extended Data Fig. 8e,f). Thus, budding yeast and other species utilise different elements to tightly tether the Pol III heterotrimer to the polymerase core so that Pol III initiates transcription efficiently.

**Fig. 3.**
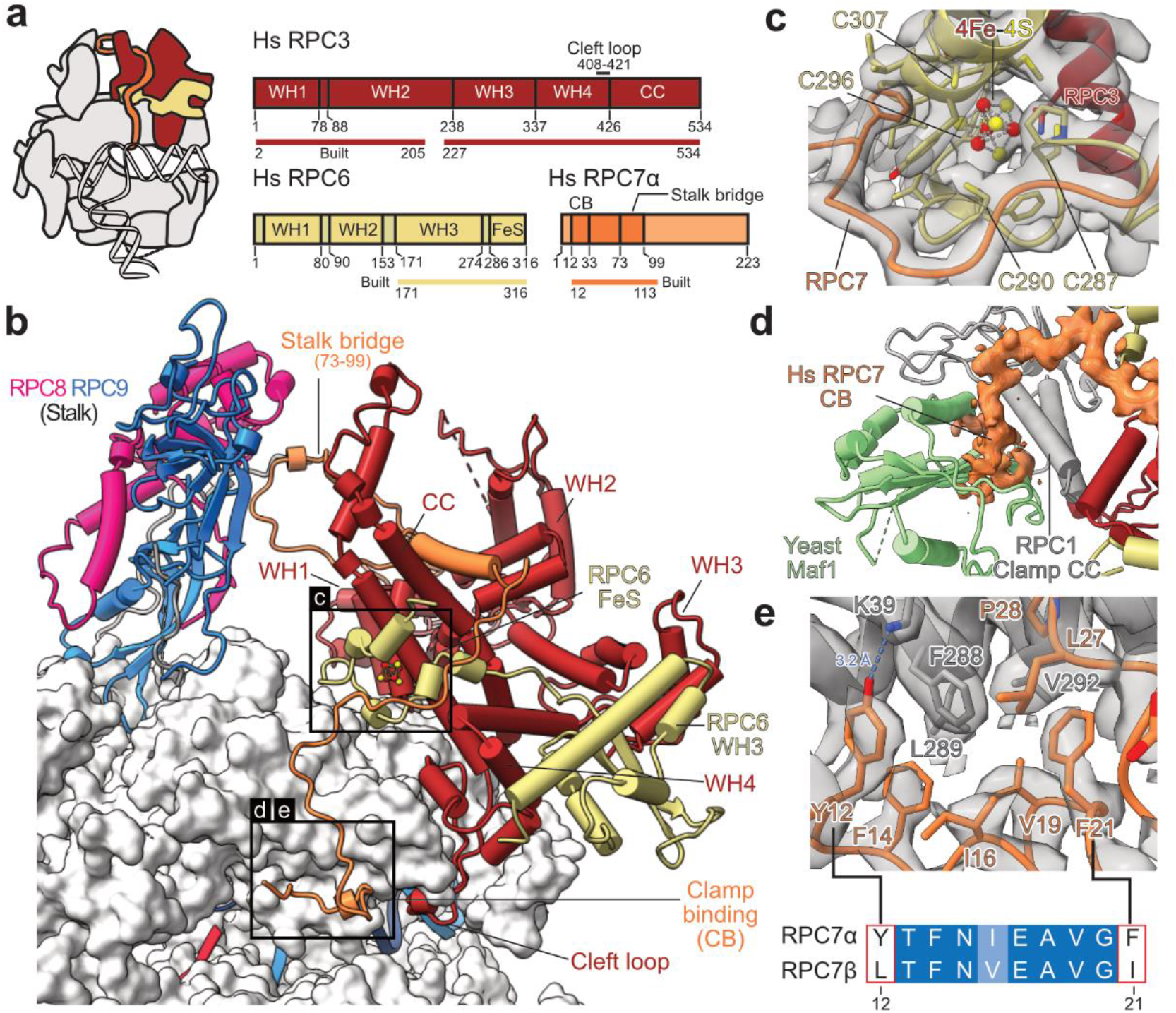
Architecture of human RPC3-RPC6-RPC7 heterotrimer. **a**, Schematic domain organisation of subunits RPC3, RPC6, and RPC7 in the elongating human Pol III complex. Coloured bars indicate modelled regions and are labelled as ‘built’. WH – winged helix, CC – coiled-coil, FeS – Iron-sulphur (4Fe-4S) binding, CB – clamp binding. **b**, Heterotrimer (RPC3-RPC6-RPC7) and stalk (RPC8-RPC9) are shown as cartoons and bind the Pol III core shown as surface. Boxed regions highlight features that are not visible in yeast Cryo-EM structures. **c**, Structure of the RPC6 FeS domain binding the cubane 4Fe-4S cluster. Residues within 6 Å of the FeS cluster are shown as sticks, and the four coordinating cysteines are labelled. Cryo-EM density is shown as a transparent surface. **d**, Superimposition of the apo human Pol III structure with the yeast Pol III-Maf1 structure (PDB 6tut). Only Maf1 is shown and is coloured light green. Cryo-EM density of RPC7 is shown as an orange surface. **e**, Interface between the RPC1 clamp coiled-coils (CC, grey) and RPC7 (orange). RPC1 F288 is surrounded by aromatic residues of RPC7, of which Y12 and F21 are exclusive to the tumour-associated RPC7α isoform as shown by the sequence alignment between RPC7α (UniProt: O15318) and RPC7β (UniProt: Q9BT43) placed below. Corresponding Cryo-EM density is shown as a grey transparent surface. The blue dashed line indicates a putative hydrogen bond between Y12 (RPC7) and K39 (RPC1).

### RPC7 bridges heterotrimer and stalk

The structures of human Pol III further provide new insights into the function of RPC7 (C31 in *S. cerevisiae*), which is the third subunit of the heterotrimer and has been proposed to bridge the heterotrimer and the mobile stalk element of Pol III^15,16^. Improved densities for RPC7, compared to the yeast cryo-EM maps, allowed us to assign the connection between heterotrimer and stalk with high confidence (Extended Data Fig. 5e). The RPC7 stalk bridge comprises residues Y73-W99 and protrudes from a central helix that is tightly enclosed by the RPC3 coiled-coil (CC) domain and the RPC3 WH2 domain (Fig. 3b). RPC7 is anchored to the Pol III stalk with a conserved tyrosine (Y87) (Supplementary Figure 1) that forms a cation-π interaction with R107 of RPC8 and then folds back to bind the RPC3 WH2 domain (Fig. 3b, Extended Data Fig. 5e). While in yeast, the bridge was reported to only form in the process of transcription initiation^16^, we observe that human RPC7 bridges heterotrimer and stalk in both apo and elongating Pol III (Extended Data Fig. 9). The RPC7 stalk bridge moves together with the stalk upon transition from apo to elongating Pol III, despite an overall movement of the heterotrimer and stalk in opposite directions (Extended Data Fig. 9). Thus, RPC7 ensures that the connection between stalk and heterotrimer is maintained, and the main function of the RPC3 CC and WH2 domains, most likely, is to tether RPC7 to the heterotrimer.

### Tumour-associated RPC7α binds the clamp

A better understanding of RPC7 is of biomedical relevance as this subunit occurs in two isoforms: RPC7α (also known as RPC32α and POLR3G) and RPC7β (also known as RPC32β and POLR3GL). RPC7α is enriched in embryonic stem cells and cancer cells^14,34–36^, and its overexpression augments tumour transformation^14^. Depleting RPC7α also suppresses viability of prostate cancer cells^36^. Our cryo-EM structures show that the N-terminus of RPC7 (residue 10-33) runs along the RPC1 clamp domain and docks onto the RPC1 clamp coiled-coil (CC) helices (Fig. 3b,d,e). Thereby, this clamp-binding (CB) region would overlap with the Pol III repressor Maf1 in the recently reported cryo-EM structure of *S. cerevisiae* Pol III bound by Maf1^19^ (Fig. 3d). Thus, RPC7, most likely, must be replaced to enable Pol III repression by the tumour suppressor MAF1 (the homolog to yeast Maf1). In the Pol III-Maf1 structure, the interaction between Maf1 and Pol III is stabilised by an aromatic stacking between W319 (*S. cerevisiae* Maf1) and W294 (*S. cerevisiae* RPC1/C160). Importantly, we observe that RPC7 associates with human RPC1 at the same interface via a hydrophobic surface that encloses F288, the human counterpart to W294 in *S. cerevisiae* RPC1/C160 (Fig. 3e). In our structure, RPC1 F288 is juxtaposed by RPC7α Y12 and potentially forms an aromatic stacking interaction and a hydrogen bond to RPC1 K39. Neither the aromatic stacking nor the hydrogen bond are possible with the corresponding RPC7β residue (L12) (Fig. 3e) suggesting that RPC7α could bind the clamp more tightly than RPC7β. In support of this hypothesis, the corresponding first 25 N-terminal residues in *S. cerevisiae* RPC7/C31 are more similar to RPC7β then to RPC7α (Supplementary Figure 1) and do not bind the clamp in all available *S. cerevisiae* Pol III structures^15–19^. We hypothesise that the RPC7α subunit specifically protects against Maf1-mediated Pol III inhibition in tumour cells, providing a potential link between RPC7 isoform selection and tumour transformation. Interestingly, methylation of arginines R5 and R9 in *S. cerevisiae* RPC7/C31 by Hmt1 increases the affinity of Maf1 to Pol III and represses Pol III activity^37^. In humans, RPC7β but not RPC7α is a substrate to Hmt1^37^, reinforcing the idea that incorporation of RPC7α results in a Pol III isoform that is resistant to Pol III inhibition by Maf1.

### Disease-associated mutations of Pol III

Mutations in human Pol III have been linked to neurodegenerative disorders like Hypomyelinating Leukodystrophy (HLD) affecting white matter of the central nervous system^38–44^, developmental disorders such as Wiedemann-Rautenstrauch Syndrome (WDRTS), a progeroid neonatal disorder^45,46^ and Treatcher Collins Syndrome (TCS) affecting craniofacial development^40,47^ and susceptibility to infections by Varicella Zoster Virus (VZV), which can lead to severe central nervous system disorders and pneumonitis^48,49^ (Fig. 4a). The high-resolution cryo-EM structure of human Pol III with side-chain densities for all, except one, of the mutated residues (Supplementary Fig. 2) allows us to confidently map and investigate these mutations on a molecular level. Point mutations are found mainly in three clusters: in the core of the complex close to the active site, in subunits RPAC1 and RPAC2 shared with Pol I and in the vicinity of RPC10 where they might affect termination or re-initiation^44^ (Fig. 4b). On the contrary, the majority of VZV-causing mutations are found on the Pol III surface (Fig. 4a,b). Interestingly, VZV-related mutations have no phenotype in the absence of a viral infection^48^. Therefore, the identified mutations in RPC3 and RPC5 presumably have little effect on canonical nuclear Pol III transcription, but specifically impair cytosolic transcription of viral AT-rich DNA by Pol III to induce interferon pathways^48^ and exert control over viral replication^49^. We speculate that, in the nucleus, where transcription factor TFIIIB interacts extensively with RPC3 and RPC5 to facilitate specific transcription initiation, the effect of these mutations might be buffered, whereas in the cytosol, transcription initiates unspecifically and the identified mutations impair Pol III activity.

**Fig. 4.**
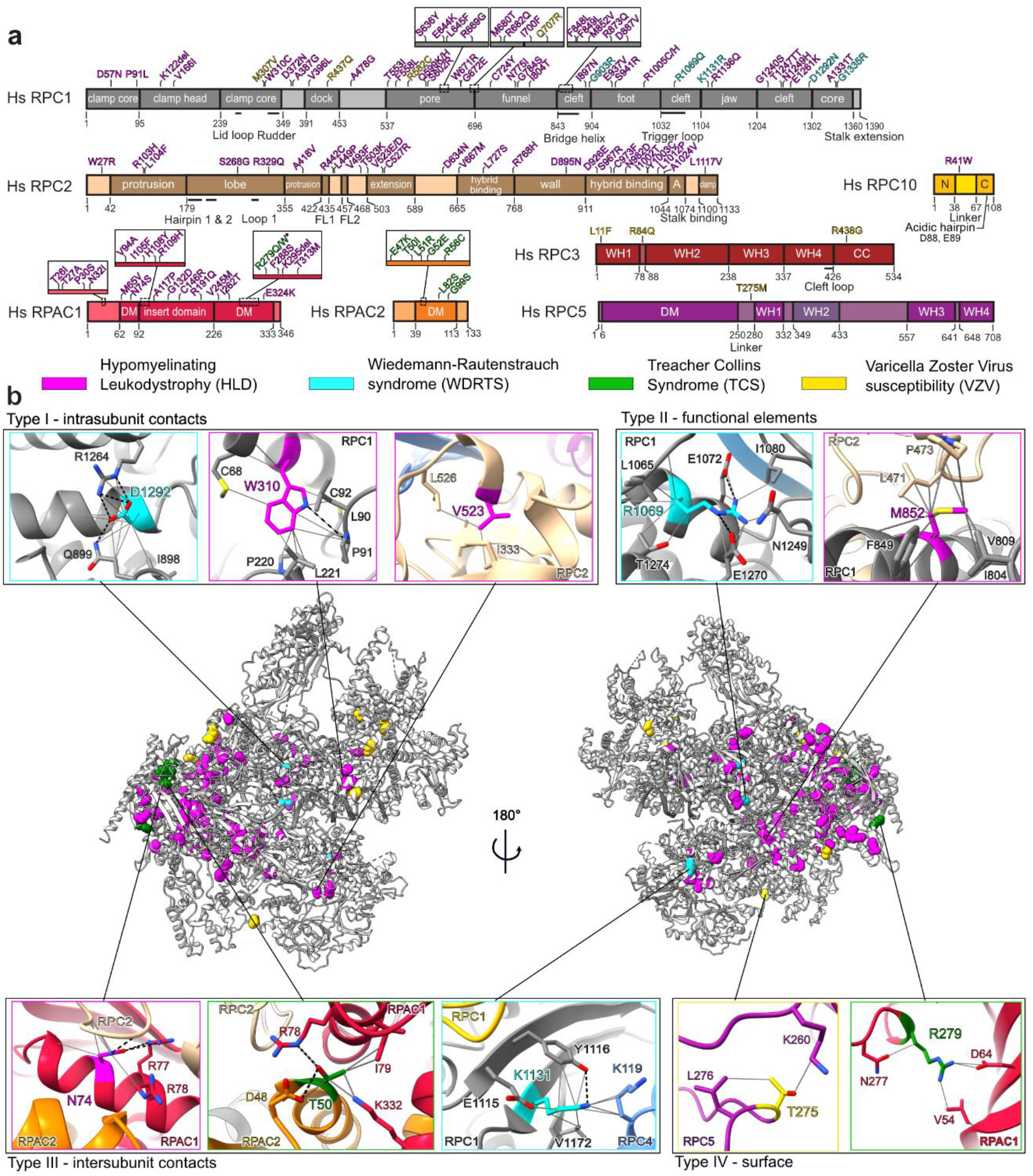
Disease-associated mutations of Pol III. **a,** Schematic representation of Pol III subunits affected by disease-associated mutations mapped onto the domains. R279Q/W mutations (starred) associated with TCS^47^ was also reported in individuals with HLD^40^. A – anchor; N – N-terminal domain; C – C-terminal domain; DM – dimerization domain; WH – winged-helix; CC – coiled-coil. **b**, Disease-associated mutations (coloured spheres associated with each disease) mapped on a cartoon representation of human Pol III. Close up views of representative mutations belonging to one of 4 identified types (see Methods): I – intrasubunit contacts, II – functional elements, III – intersubunit contacts and IV – surface are shown. Grey solid lines – contacts between residues, black dotted lines – hydrogen bonds.

We used 110 point mutations found through genetic studies of patients (Fig. 4a, Supplementary Table 2 and references therein) and classified them into four types explaining their likely mode of function on a molecular level (Supplementary Table 2). Type I mutations disturb the core of a given subunit (Fig. 4b, Supplementary Table 2). Type II mutations affect functional elements such as the bridge helix (F848L, F849L, M852V, R873Q and D887V mutations linked to HLD) or the trigger loop (R1069Q mutation linked to WDRTS) (Fig. 4a,b). Type III mutations are located at the interface between subunits and can impair the assembly of the complex, as observed in the case of the N32I and N74S mutations (Fig. 4b) linked to HLD^40^. Type IV mutations comprise residues making few contacts and being surface exposed, and we could observe that the majority of them link to VZV (Extended Data Fig. 10). This classification can help to explain the severity of disease phenotypes linked to a given mutation. In case of TCS, mutations found in RPAC2 are dominant (and thus need to be highly deleterious)^47^ and are all classified as type I or III, while the only recessive TCS mutation R279Q/W found in RPAC1^40,47^, which we assigned to both type I and IV, is associated with less severe structure disturbance (Fig.4b, Extended Data Fig. 10b,c). In the case of HLD, the subunit in which the mutation is found often correlates with the severity of a disorder. HLD patients having mutations mapping to RPAC1 experience the most severe symptoms, followed by those with mutations in RPC1^42,43^. Many of the type III mutations map to RPAC1 impairing biogenesis of Pol III^40^, while the majority of the type II mutations map to RPC1 (Extended Data Fig. 10c).

Additional parameters to assess the impact of pathogenic mutations are the calculated solvent accessible surface area (SASA) and the residue degree, which quantifies the number of its contacts to other residues (Supplementary Table 2). We find that higher SASA and a lower degree of mutated residues link to a less severe clinical phenotype. For instance, the WDRTS mutation K1131R (SASA: 19.8 Å^2^, degree: 17) is considered to be less deleterious than R1069Q (SASA: 10.5 Å^2^, degree: 23) and D1292N (SASA: 8.4 Å^2^, degree: 21)^45^. Accordingly, the human Pol III structure is a useful resource for investigating the molecular environment of mutations and together with a comprehensive point mutation report (Supplementary Table 2) will advance the understanding of the genotype-phenotype relationship in various diseases.

### Concluding remarks

The cryo-EM structures of human Pol III promote our understanding of the Pol III transcription machinery and provide molecular insights into how Pol III misregulation is linked to human pathogenesis. Accordingly, we provide structure-based explanations of how subunit RPC10 functions in transcription termination and how cancer cells may specifically utilise isoform RPC7α to uncouple Pol III activity from repression by the tumour suppressor MAF1. The structure determination of human Pol III was facilitated by CRISPR-Cas9 mediated gene-editing of human suspension cells that enabled purification of homogenous and intact protein sample. The possibility to produce human Pol III in sufficient amounts for functional analysis and high-resolution structure determination may also facilitate the development, characterisation and structure-guided optimisation of inhibitors specifically targeting upregulated Pol III in human cancer cells. We also envision that CRISPR-Cas9 approaches will be increasingly used to produce other challenging human protein complexes to gain structural, functional and biomedical insights.

## ACKNOWLEDGEMENTS

We thank F. Dossin for giving valuable advice on CRISPR-Cas9 and sharing a plasmid used in gene editing, F. Weis and W.J.H. Hagen (EMBL Cryo-Electron Microscopy Service Platform) for EM support, P. Haberkant and M. Rettel (EMBL Proteomics Core Facility) for carrying out MS analysis, T. Hoffmann and J. Pecar for setting-up and maintaining the high-performance computing environment and current members of the Müller lab and S. Eustermann for discussions. M.G. was supported by a Boehringer Ingelheim Fonds PhD fellowship. M.G, A.D.M and A.L. acknowledge support by the EMBL International PhD program.

## AUTHOR CONTRIBUTIONS

C.W.M. initiated and supervised the project. M.G. initiated the project, carried out EM grid preparation, data collection and processing, model building and interpreted the structures. A.D.M. generated HEK293F cell lines via CRISPR-Cas9 mediated gene-editing, purified proteins and analysed Pol III mutations, M.K.V. assisted with structural model building, interpretation of the structures and advices with EM data processing. A.L. and A.B. performed the phylogenomic analysis. F.B. did RNA extension and cleavage assays. H.G. performed cell culture work, advised and assisted with CRISPR-Cas9 mediated gene-editing and protein purification. M.G., A.D.M. and C.W.M. wrote the manuscript with input from the other authors.

## AUTHOR INFORMATION

The authors declare no competing financial interests. Correspondence and requests for materials should be addressed to C.W.M.

## METHODS

### Generation of a cell line with endogenously tagged Pol III

CRISPR-Cas9 methodology was used to generate a stable, homozygous cell line with tagged Pol III. HEK293F cells were engineered based on a modified general protocol^50^. A 20 bp gRNA (5’-ACTGAGCTTGGATGCTTCTG-3’) was designed using the online tool crispor.tefor.net and cloned into the BpiI site of pSpCas9(BB)-2A-GFP (PX458), a gift from Feng Zhang (Addgene plasmid # 48138)^51^. The donor sequence with 700 bp homology arms on each side of the insert site was designed based on the HEK293 reference genome^52^ and synthesised by GenScript into pUC57-Mini plasmid. A tag with mCherry, Strep II, 6xHis and P2A peptide^53^ followed by a blasticidin resistance gene and the SV40 termination signal was introduced to the C-terminus of the RPAC1 subunit. HEK293F cells cultured in DMEM medium supplemented with 10% FBS and 1% L-glutamine were transfected with the donor plasmid and the plasmid pSpCas9(BB)-2A-GFP (PX458) containing the gRNA, using polyethylenimine reagent in Opti-MEM I Reduced Serum Medium (Gibco). 5 days post-transfection cells were selected with 5 μg/mL blasticidin (Thermo Fisher Scientific). When the survival rate of the cells improved, they were sparsely seeded to form colonies from single cells. 48 hours after seeding, single clone colonies were picked by pipetting into 96 well plates and genotyped. To check for the insertion of the tag one primer annealing within the insert and one annealing within a homology arm were used for PCR. Clones homozygous for the insert were selected and expanded. Correct insertion was validated by Western blotting against mCherry (antibody ab167453; Abcam) and PCR on extracted DNA (GenElute Mammalian Genomic DNA Miniprep Kit; Sigma-Aldrich). Selected clones were finally adapted to growth in suspension in Expi293 Expression Medium (Thermo Fisher Scientific).

### Protein purification

Cells were grown up to 7 x 10^6^ cells/mL in Expi293 medium, harvested by centrifugation, washed in PBS and flash-frozen in liquid nitrogen for storage before protein purification. Cell pellets were resuspended in lysis buffer containing 25 mM HEPES pH 7.5, 150 mM (NH_4_)_2_SO_4_, 5 mM MgCl_2_, 5% glycerol, 20 mM imidazole, 0.5% Triton-X100, 2 mM β-mercaptoethanol in the presence of EDTA-free protease inhibitor cocktail (Roche) and benzonase (Sigma-Aldrich). Cells were sonicated and centrifuged at 235,000 g for 1 h at 4 °C. After filtration, the lysate was loaded on a 5 mL HisTrap HP column (GE Healthcare) washed with Ni wash buffer (25 mM HEPES pH 7.5, 150 mM (NH_4_)_2_SO_4_, 5 mM MgCl_2_, 5% glycerol, 20 mM imidazole, 2 mM β-mercaptoethanol) and eluted with buffer containing 500 mM imidazole. The elute was then incubated for 1 h at 4 °C with Strep-Tactin beads (IBA Lifesciences) before application to a gravity column. After washing with Strep wash buffer (25 mM HEPES pH 7.5, 150 mM (NH_4_)_2_SO_4_, 5 mM MgCl_2_, 5% glycerol, 2 mM β-mercaptoethanol), the protein was eluted with buffer containing 100 mM (NH_4_)_2_SO_4_ and 20 mM biotin. The recovered complex was subsequently purified by ion exchange on a MonoQ column (GE Healthcare). Pol III eluted at about 700 mM (NH_4_)_2_SO_4_. It was concentrated on 100K spin concentrators (Merck Millipore) to about 3 mg/mL and then buffer exchanged to EM buffer (15 mM HEPES, pH 7.5, 150 mM (NH_4_)_2_SO_4_, 5 mM MgCl_2_ and 10 mM DTT). The purity of the sample was assessed by SDS-PAGE and Coomassie blue staining, and the identity of subunits was confirmed by mass spectrometry. The concentrated sample was either flash-frozen in liquid nitrogen and stored at −80 °C for RNA extension assays or used directly for cryo-EM sample preparation. Yeast RNA Pol III was purified as previously described^54^.

### Transcription scaffold preparation

The following nucleic acid oligonucleotides (Sigma-Aldrich, HPLC-grade) were used for the transcription scaffold containing a 12-nt mismatch that was freshly assembled prior to cryo-EM sample preparation of elongating Pol III (see below). Template DNA: 5’-GTTTTAAGTGGAGCTGGATGCTCTGTGGCTCATGGCAACGACCACG-3’, non-template DNA: 5’-CGTGGTCGTTGCCATGTTAACGGACTAGTCCAGCTCCACTTAAAAC-3’, RNA: 5’-UAUGCAUAACGCCACAGAG-3’. Single-stranded template- and non-template DNA at 100 μM were mixed, incubated at 95 °C in a thermocycler (Bio-Rad) for 3 min, and the temperature was incrementally decreased to 25 °C at a rate of 1 °C/min. The double-stranded DNA (2 μL) was supplemented with 3 μL of hybridisation buffer (40 mM HEPES pH 7.5, 24 mM MgCl_2_, 200 mM NaCl, 20 mM DTT) and mixed with 1 μL of RNA (100 μM), which was pre-heated in a thermoblock for 3 min at 55 °C. The DNA-RNA mixture was incubated at 45 °C for 3 minutes, cooled down to 20 °C at a rate of 0.7 °C/min, and the annealed DNA-RNA transcription scaffold was kept on ice until usage.

### RNA extension and cleavage assays

For RNA extension assays, the RNA primer (see above) was first labelled at its 5’ end using [γ-^32^P]ATP and T4 PNK. After PAGE purification, the radioactive RNA primer was assembled together with template and non-template strands (see above) in 20 mM HEPES pH 7.5, 100 mM NaCl and 6 mM MgCl_2_ and 10 mM DTT as indicated above. To assemble the reaction mixture, 2 pmol of the DNA/RNA-^32^P were incubated with 4 pmol of human or yeast RNA Pol III for 10 min at room temperature, before addition of 0.2 mM of NTPs as indicated in 20 mM HEPES pH 7.5, 60 mM (NH_4_)_2_SO_4_, 10 mM MgSO_4_ and 10 mM DTT, and further incubated for 45 min at 37 °C. The reaction was stopped by addition of 8M Urea/20% saccharose. The samples were heated for 3 min at 95 °C before loading on denaturing 17% PAGE. The radioactive products were recorded using phosphor-imaging screens (Fujifilm) for capturing digital images. For RNA cleavage assays, the procedure was the same except that no NTPs were added. The incubation times were 0, 5 min, 20 min, 60 min and 120 min.

### Negative stain EM

Negative stain EM was used to assess sample quality of purified human Pol III. For the preparation of negative stain grids, human Pol III was diluted to a concentration of 50 nM using EM buffer, and 3.5 μL sample were applied to carbon-coated 400 mesh copper grids (Plano) that were glow discharged using a Pelco EasyGlow instrument. After 60 s incubation, grids were washed twice with EM buffer, stained with 1% (w/v) uranyl acetate staining solution and air dried. Negative stain EM data were acquired on a Tecnai T12 transmission electron microscope (TEM) operated at 120 keV (FEI) with a 4k x 4k CCD camera. Images were recorded at −1 μm defocus and at a magnification of 68,000x corresponding to a pixel size of 1.6 Å/pixels. A dose rate of 40 e/Å^2^ was used during image acquisition.

### Cryo-EM sample preparation

Freshly purified human RNA Pol III was used for cryo-EM sample preparation. Grids were prepared using a Vitrobot Mark IV (Thermo Fisher Scientific), which was set to 100% humidity and 4 °C. The following blotting parameters were used for both copper and gold grids: wait time 10 s, blot force 4, blot time 4 s. For apo RNA Pol III, 3 μL of protein at 1.2 mg/mL in EM buffer was supplemented with 4 mM CHAPSO (final concentration), applied to plasma-cleaned 200-mesh Cu R2/1 or 300-mesh Au R1.2/1.3 grids (both Quantifoil) and plunge-frozen in liquid ethane. For elongating Pol III, the complex at 2 mg/mL in EM buffer was mixed with an equimolar amount of the transcription scaffold, incubated for 60 min on ice, supplemented with 4 mM CHAPSO (final concentration), applied to 200-mesh Cu R2/1 grids and plunge-frozen in liquid ethane as described above. The cryo-EM grids were plasma-cleaned for 30 s using a NanoClean plasma cleaner (Fischione Instruments, Model 1070) and a mixture of 75% argon and 25% oxygen before usage.

### Cryo-EM data collection and processing

Automated data acquisition was performed using SerialEM^55^. Grids were screened using a Talos Arctica TEM operated at 200 keV (Thermo Fisher Scientific) and equipped with a Falcon III direct electron detector (Thermo Fisher Scientific). A small dataset of Apo Pol III applied to a 300-mesh Au R1.2/1.3 grid (Quantifoil) was recorded at a magnification of 92,000x corresponding to a pixel size of 1.566 Å/pixel. A total of 753 image stacks (12 frames, exposure time 2.1 s) were collected using a dose rate of 3.91 e/Å^2^/frame (46.92 e/Å^2^ total dose) at a defocus range of 1-3.5 μm on an FEI Talos Arctica microscope equipped with a Falcon III detector operated in linear mode. High-resolution cryo-EM data on apo Pol III, plunge frozen on a 200-mesh Cu R2/1 grid (Quantifoil), was collected on a Titan Krios TEM operated at 300 keV (Thermo Fisher Scientific) and equipped with a Quantum energy filter (Gatan) and a K3 direct detector (Gatan). A magnification of 105,000x was used, corresponding to a pixel size of 0.822 Å/pixel. We recorded 9,544 image stacks (40 frames, exposure time 1.2 s) in counting mode using a dose of 0.95 e/Å^2^/frame (38.11 e/Å^2^ total dose) and a defocus range of 0.75-2.25. High-resolution cryo-EM data on elongating Pol III was collected on a 300 keV Titan Krios TEM equipped with a K2 direct detector (Gatan). We used a magnification of 130,000x, corresponding to a pixel size of 1.05 Å/pixel, and collected 9,488 micrographs (40 frames, exposure time 11.2 s) in counting mode at a dose of 1.01 e/Å^2^/frame (40.35 e/Å^2^ total dose). The defocus range was 0.75-2.00 μm.

For all datasets, initial frame alignment, contrast transfer function (CTF)-estimation, dose-weighting and automated particle picking was performed using Warp^56^. The images were further processed with RELION 3.1^57^. In order to run particle polishing at later stages during processing, the micrographs, acquired with a Titan Krios, were also frame-aligned and dose-weighted using RELION’s own implementation of the MotionCor2-algorithm^58^ and CTF-corrected using Gctf^59^.

For the Talos Arctica dataset, a total of 65,009 particles were picked and extracted with a box size of 230 pixels. The images were processed with cryoSPARCv2^60^. After 2D classification, 48,473 particles were chosen to generate an initial model using 2 classes. One of the initial models resembled the shape of yeast RNA Pol III and was chosen for homogenous refinement using 37,932 particles. The resulting map of apo human Pol III reached a resolution of 8.0 Å.

In case of the high-resolution apo Pol III dataset, 304,683 particles were picked and extracted with a box size of 328 pixels using Warp. Subsequently, global 3D classification in RELION was performed using 4 classes. As a reference model, we used the 8.0 Å apo human Pol III map obtained from the Talos Arctica dataset, and low-pass filtered it to 60 Å. A single class was selected, and the 111,289 particles were subjected to 3D refinement yielding a 3D reconstruction at 3.8 Å resolution. The same set of particles was re-extracted with a 440 pixels box size from the micrographs that were pre-processed in RELION and subjected to 3D refinement. This was followed by two iterative rounds of particle polishing and CTF refinement, which included the estimation of anisotropic magnification and high-order aberrations^61^. Subsequent 3D refinement and map sharpening yielded a 3D reconstruction at 3.3 Å resolution (Map A), which was further subjected to multi-body refinement^62^ using two masks. Mask 1 covered the core of the polymerase and subunits RPC4/5/10, and mask 2 included the stalk, the clamp and the RPC3/6/7 heterotrimer. The two resulting maps, A1 and A2, were refined to 3.2 and 3.4 Å resolution, respectively, and used for model building of the apo Pol III structure.

For the elongating Pol III complex, 367,717 particles, extracted with a box size of 220 pixels, were subjected to 3D classification using a 60 Å low-pass filtered reconstruction of apo human Pol III as a reference map. One of the 4 classes, containing 166,071 particles, showed clear density for the bound downstream DNA and was refined to a resolution of 3.2 Å. After particle polishing and CTF refinement, the resolution was further improved to 2.8 Å (Map B). To improve map quality of less well-resolved regions, we performed masked classification in RELION without image alignment and asked for 2 classes per classification procedure. Soft masks were created using the molmap command in UCSF Chimera^63^ on the regions of interest followed by mask creation in RELION.

First, we applied a mask covering the downstream DNA and the DNA/RNA hybrid. Using a regularization parameter T of 10 during 3D classification, we obtained a class containing 70,492 particles, which showed improved signal corresponding to the DNA/RNA hybrid in the active site (map C, 3.1 Å). During model building (see below), we observed additional density starting from the linker domain of RPC10 and extending into the polymerase funnel. Therefore, we placed the C-ribbon domain of RPC10 into the funnel using Chimera, created a soft mask and performed masked classification on this area using a regularization parameter of 20. Class 1 contained 30,525 particles and showed clear density inside the funnel (map D, 3.1 Å) and, compared to map B, only little residual density corresponding to the RPC10 C-ribbon domain outside the funnel. Class 2, containing 100,152 particles, showed no density inside the funnel but stronger signal belonging to the RPC10 ribbon outside the funnel (map E, 2.9 Å). At lower threshold levels, we observed additional density close the cryo-EM density that corresponds to the RPC4/5 heterodimer orthologue C53/C37 in *S. cerevisiae*, leading us to assume that this density belongs to the additional RPC5 WH domains. Thus, we first performed masked classification using a mask that covers the RPC4/5 dimerization module, followed by masked classification on the additional density. We manually place a homology model obtained for the WH1 domain of RPC5 (278-352), created using the Phyre^64^ web server, into this density. The fit of the placement was also inspected using COOT^65,66^, and the C-terminal helix (339-352) of the homology model was re-arranged so that it fitted a close-by density that resembled the shape of a longer helix. The fitted homology model was used for mask creation, and masked 3D classification was performed using a regularization parameter T of 40. One class contained 21,861 particles and showed improved density corresponding to the RPC5 WH 1-2 domains binding the polymerase core (map F, 3.3 Å).

We noticed that mostly the polymerase clamp, stalk and heterotrimer change conformation during the transition from apo to elongating Pol III whereas the remaining parts stay rigid. Therefore, we merged the two datasets using RELION in order to improve densities for less well-resolved regions. Pol III apo particles (map A) were re-extracted, binned to a pixel size of 1.05 Å/pixel, merged with the Pol III EC particles (map B) using RELION, and imported into cryoSPARC for further processing steps. In cryoSPARC, the particles were subjected to homogenous refinement, followed by CTF refinement and non-uniform refinement^67^. The resulting 3D reconstruction (map G, 2.9 Å) showed improved density for an area near the RPC4/5 heterodimer that extended towards the polymerase jaw domain. The merged dataset was also used to repeat masked classification on the RPC5 C-terminal extension to resolve a second conformation of the WH domains. We followed the same strategy as described for map F but, this time, focused on the counterpart map, in which additional density was visible at lower threshold levels. A second round of masked classification on the RPC5 WH domains yielded a map containing 31,737 particles that showed density for the WH domains in the alternative conformation. The corresponding density could be slightly improved by non-uniform refinement in cryoSPARC (map H, 3.4 Å). Resolution values reported above are based on the gold-standard Fourier shell correlation (FSC) using the 0.143 cut-off criterion^68^ and were obtained using RELION post-processing. Local resolutions of map A to map F were estimated using the local resolution estimation tool implemented in RELION. For map G and H, MonoRes^69^ was used to estimate the local resolution range.

To get insights into conformational dynamics, we used the 3D variability analysis (3DVA)^26^ tool implemented into cryoSPARC. Particles belonging to map B, obtained via processing in RELION, were imported into cryoSPARC, subjected to homogenous refinement followed by 3DVA using a soft mask covering the complete map.

### Structural model building and refinement

An initial model of apo human RNA Pol III was constructed using homology models and available structures. Homology models of RPC1, RPC2, RPC4 and RPAC2 were obtained from the SWISS-MODEL Repository^70^. Homology models of RPC5, RPC6, RPC7, RPC8, RPC9, RPC10 and RPAC1 were generated using Phyre2. The homology models and the structures of RPC3-RPC7 (PDB: 5AFQ) and of RPABC1, RPABC2, RPABC3, RPABC4, RPABC5 (PDB: 5IY6) were first aligned to their homologues counterparts in the Pol III apo structure from S. cerevisiae (PDB: 5FJ9). The constructed 17-subunit model was fitted as a rigid body into the cryo-EM density map of apo human Pol III using Chimera, and the individual polypeptide chains were fitted individually as rigid bodies followed by manual model building and real-space refinement using COOT.

For the elongating RNA Pol III structure, the model of the apo human Pol III was first fitted into the cryo-EM density of the elongating Pol III map B in Chimera, and the chains were adjusted manually in COOT. Cryo-EM density, corresponding to the upstream DNA, was observed in the DNA-binding cleft, and the DNA/RNA-duplex molecule from the elongating *S. cerevisiae* RNA Pol III structure (PDB: 5FJ8) was placed into the corresponding density. Interatomic distance restraints of the DNA/RNA molecule were generated using ProSMART^71^, implemented in the most recent version of COOT^66^, and the nucleic acids were first fitted automatically into the density and refined using the all-molecule-refine option in COOT. The nucleotide bases were then mutated to match the sequences of the used nucleotides, followed by manual fitting and refinement against map C to obtain a more accurate fit.

When building the model of RPC10, we observed that the density corresponding to the linker and the C-ribbon domain needs to be traced differently compared to the cryo-EM apo structure of *S. cerevisiae* RNA Pol III (PDB: 5fja). From residue E48 ongoing, we observed continuous density that split-up and either extended into the Pol III funnel or towards the polymerase periphery, which could be resolved by masked classification (see above). For the building of the RPC10-C-ribbon domain, we used map D (inside funnel) and map E (outside funnel). In both cases, a homology model (67-107) was placed into the corresponding densities using Chimera, followed by manual fitting and refinement using COOT. In addition, ISOLDE^72^ (implemented into the ChimeraX software^73^) was used to reduce the clash score and to optimise fitting with applied secondary-structure restraints on predicted beta-sheets based predictions obtained via the PSIPRED webserver^74^. Placement of the zinc atom in the C-ribbon domain was partially guided by the crystal structure of the C-ribbon of *S. cerevisiae* TFIIS bound to *S. cerevisiae* RNA Pol II (PDB: 3PO3).

For building the FeS domain of RPC6, we placed the coordinates of a cubane 4Fe-4S cluster (obtained from PDB: 2B3Y) manually into a strong density signal within the heterotrimer region that was even visible at very high threshold levels. The FeS binding domain could be built d*e novo* by extending the RPC6 homology model into visible density extending towards the 4Fe-4S signal.

For the C-terminal extension of RPC5, homology models of WH1 (278-332) and WH2 (349-433), generated via Phyre2, were first placed manually into the density (map F) using Chimera and manually adjusted and extended in COOT. The model of the WH domains was further placed in map H, in which the C-terminal extension adopts an alternative conformation 2. The WH1-WH2 domain was placed as a rigid body into the ‘conformation 2’ density in Chimera and flexibly fitted using the chain-refine command in COOT in two iterations, first with and second without distance restraints. Manual fitting of individual side-chains was not performed for the RPC5 conformation 2, and the model was trimmed by 25 peripheral residues. The DM extension and the jaw-binding domains of RPC4 were built using map G.

For model building, all cryo-EM density maps were, if not otherwise stated, locally sharpened using LocalDeblur^75^, integrated into the software framework Scipion^76^. At a later stage during model building, we also used the density modification tool^77^ implemented into PHENIX^78^ to further improve the map qualities.

PHENIX real-space refinement was used to refine the built models against the sharpened cryo-EM maps. All refinement steps were performed using Ramachandran- and rotamer restraints. For fine-tuning of model building, combining partial models and refinement, the following strategy was used: partial maps of elongating Pol III derived from RELION multi-body refinement (map B1, B2) were sharpened via density modification and used to fine-tune the model of elongating Pol III. These partial models were refined against map B1 and B2, combined and refined with secondary-structure and reference-model restraints, derived from the partial models, against map B. The model was subjected to ISOLDE and Ramachandran, and rotamer outliers were adjusted, following refinement against map B using and reference-model restraints. For refinement of more flexible regions, the following partial-model and map pairs were used for refinement: DNA/RNA – map C; RPC10-C-ribbon inside funnel – map D; RPC-10-C-ribbon outside funnel – map E; RPC5 C-terminal extension (conformation 1) – map F; RPC4 DM extension and jaw-binding domain – map G: RPC5 C-terminal extension (conformation 2) – map H. Model-based restraints for partial models were applied when refining against other maps. For the complete elongating Pol III model, partial models were combined and refined against map B using reference-model restraints. Two disease-associated residues (RPC1 W310, RPC2 W27) were manually adjusted to better fit the cryo-EM density without an additional real-space refinement step in PHENIX, and the resulting final elongating Pol III model (EC-1 Pol III) was subjected to the PHENIX comprehensive validation tool to obtain final refinement statistics. For the apo Pol III model, nucleic acids and the RPC5 C-terminal extension of elongating Pol III model were removed, and chains were manually readjusted to fit the density of apo Pol III (map A), and the model was refined against map A using reference-model restraints.

### Phylogenomic analysis of Pol III RPC5 subunits and basal transcription factors

The phylogenetic tree of Eukaryotic species was obtained from the NCBI Taxonomy Database^79^. Reference proteomes for selected species were downloaded from the UniProt website^80^ and are listed in Supplementary Table 1. In *Zea mays*, the RPC5 homolog in the representative proteome (UniProt: A0A1D6LJU7) was a partial 340 residue protein with part of the N-terminal missing and was replaced by the UniParc protein UPI0002207C81, a full 677 residue version of the RPC5 protein. Homologs for proteins of interest in this study were retrieved using the HMMER tool^81^, using Pfam^82^ domain definitions whenever possible to improve the reliability of the hits. The following UniProt and Pfam identifiers were used for homology searches: SNAPc5 (UniProt: O75971, Pfam: PF15497), SNAPc2 (UniProt: Q13487, Pfam: PF11035), SNAPc4 (UniProt: Q5SXM2), SNAPc3 (UniProt: Q92966, Pfam: PF12251), SNAPc1 (UniProt: Q16533, Pfam: PF09808), Brf1 (UniProt: Q92994, Pfam: PF07741), RPC6 (UniProt: Q9H1D9, Pfam PF05158), Brf2 (UniProt: Q9HAW0), RPC5 (UniProt: Q9NVU0, Pfam PF04801). Furthermore, Pfam family Sin_N has been renamed to RPC5, TFIIB has been improved to cover domains in Brf2, and SNAPc19 family has been iterated to include further remote homologs. Individual domains of RPC5 protein (DM and WH1-4) were assigned based on a multiple sequence alignment of all the RPC5 homologs. The Interactive Tree of Life (iTOL) online tool^83^ was used to create the final annotated phylogenetic tree.

### Analysis of Pol III mutations

Residues found to be affected by point mutations in human genetic studies were visualised using ChimeraX. Contacts and hydrogen bonds at 0.4 Å were considered. Contacts were also inspected using Protein Contacts Atlas^84^, and it was used to create Cytoscape^85^ complex network. Degree of each residue was calculated using igraph^86^ using the degree function. Solvent accessible surface area (SASA) was calculated with FreeSASA^87^. The type of mutation was assigned based on visual inspection, location within the structural elements (for type II), contacts to residues from other subunits (for type III) and SASA (above average for type IV). Residue degree was used to access the potential severity of mutation with higher degree giving a likely more deleterious outcome of a mutation.

### Data availability

Cryo-EM maps of human Pol III (map A to map G) have been deposited to the Electron Microscopy Data Base (EMDB) under accession codes EMD-XXXX. Atomic models of apo human Pol III, elongating human Pol III EC-1, EC-2 and EC-3 have been deposited to the Protein Data Bank under accession codes XXXX. The code to perform phylogenomic analysis of Pol III subunits and basal transcription factors is available from the corresponding author upon request.

## EXTENDED DATA FIGURES AND TABLES

**Extended Data Fig. 1.**
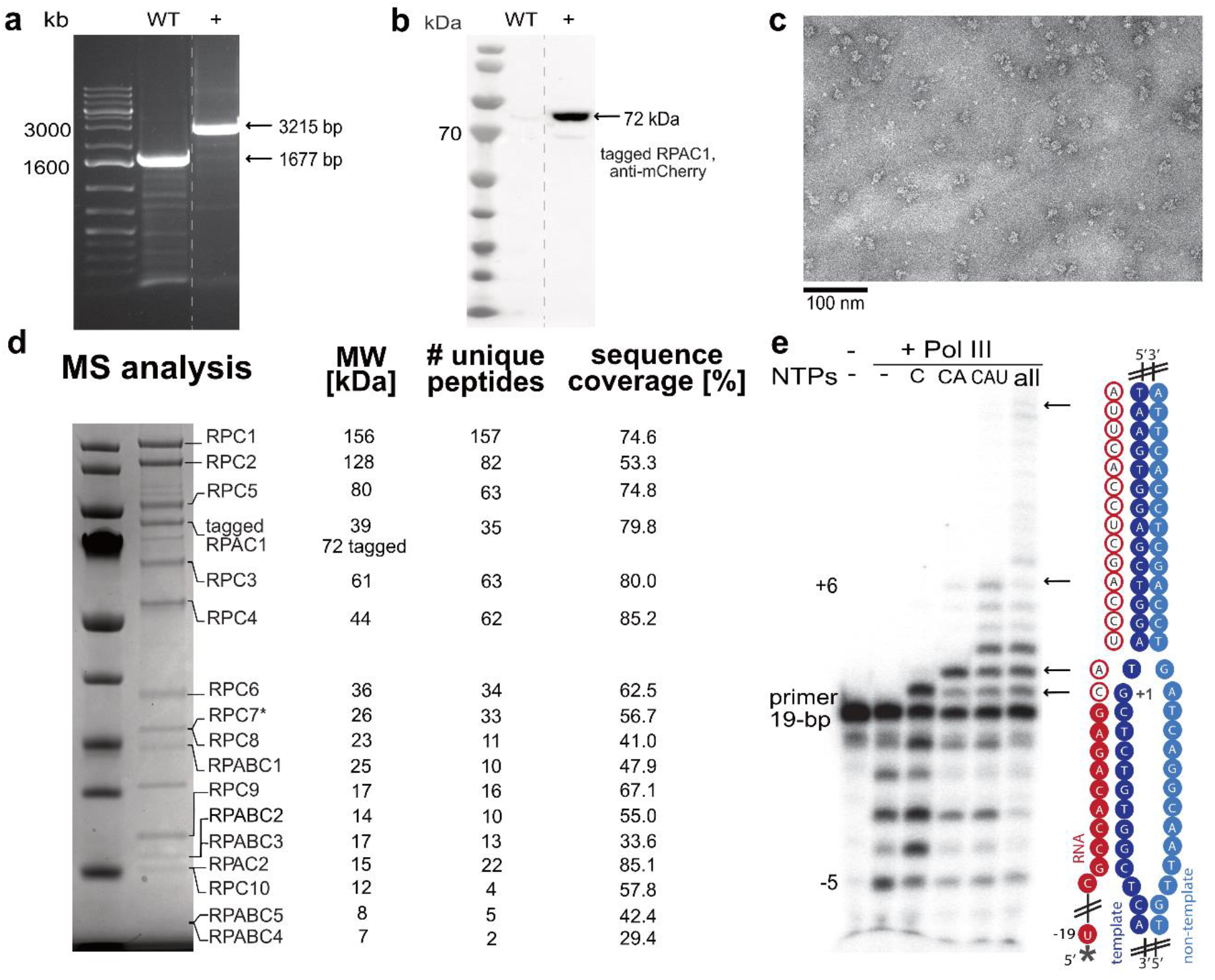
Endogenous tagging of RPAC1 and analysis of purified Pol III. **a**, Zygosity PCR with purified genomic DNA from WT cells HEK293T and produced RPAC1-Strep II-mCherry tagged cell line. The experiment was performed twice. **b**, Western blot of cell lysates from WT cells and positive for the insert cell line probed with anti-mCherry antibody (Abcam). The experiment was performed once. **c**, Representative negative stain micrograph showing homogenous sample. **d**, Coomassie-stained SDS-PAGE of purified Pol III complex. Identities of the labelled bands were confirmed by mass spectrometry with the highest scoring hit shown. For RPC7 (starred) the α isoform was detected as most abundant, while β isoform was also present in the sample. **e**, Primer extension assay confirming the activity of the purified complex. 5’ radioactively tagged RNA primer (starred) assembled with the DNA scaffold (cartoon representation) was used. Arrows indicate the extended primer products. For experimental details, see Methods. All images have been cropped for clarity.

**Extended Data Fig. 2.**
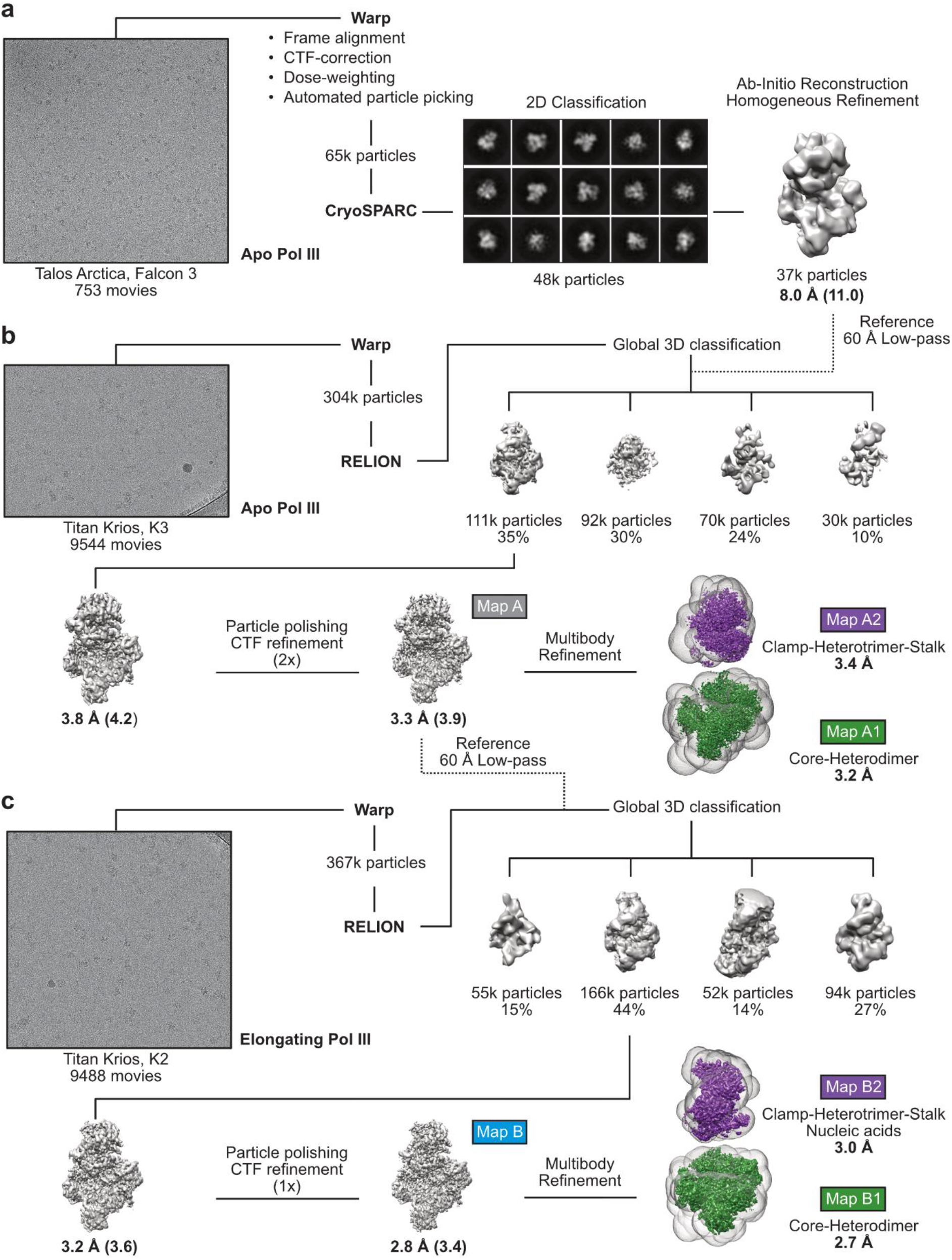
Global Cryo-EM data processing strategy of apo and elongating human Pol III. **a**, Talos Arctica screening dataset, from which a low-resolution 3D reconstruction of apo human Pol III at 8 Å was obtained. **b**, Processing pipeline of the Titan Krios data set of apo human Pol III. For global 3D classification, the low-resolution Apo Pol III map (filtered to 60 Å) was used as a reference model. **c**, Processing pipeline of the Titan Krios data set of elongating human Pol III. For global 3D classification, the high-resolution Apo Pol III map (filtered to 60 Å) was used as a reference model. High-resolution cryo-EM maps were further subjected to RELION multi-body refinement^62^ with applied masks (transparent surfaces) covering the core and heterodimer (green maps A1, B1) and the clamp, heterotrimer and stalk domains (purple maps A2, B2). Representative micrographs of the three datasets are shown on the left. Shown are the unsharpened cryo-EM maps whereas reported resolution values correspond to automatically B-factor sharpened maps obtained via RELION post-processing. Resolution values of the unsharpened maps are added with parenthesis. Shown particle numbers were rounded down. All maps, marked with coloured boxes, were used for structural model building.

**Extended Data Fig. 3.**
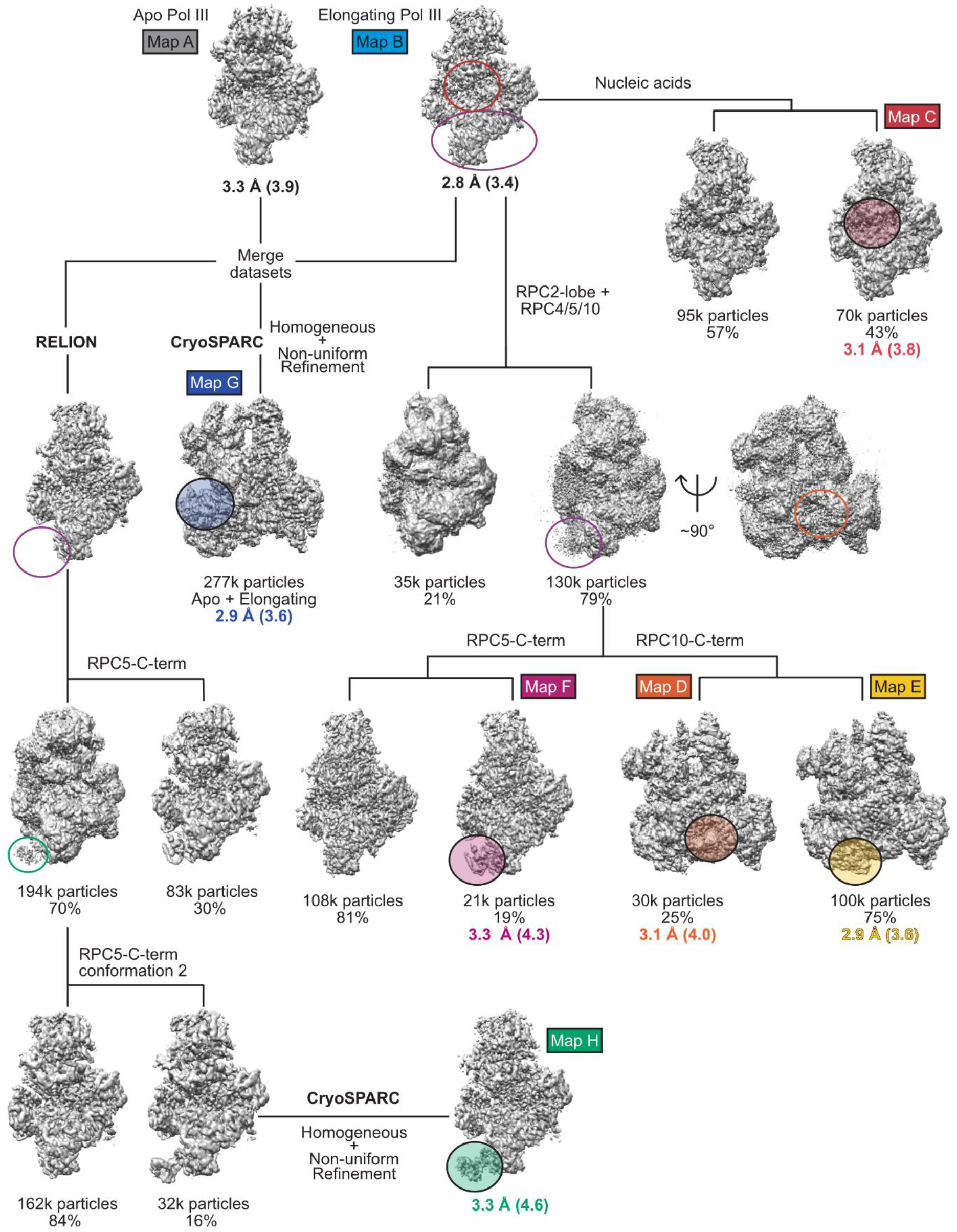
Refined cryo-EM data processing strategy to improve signals of flexible regions. Conformation heterogeneity was resolved using masked classification in RELION. Coloured unfilled circles mark regions with applied masks. Filled, transparent circles highlight elements with improved cryo-EM density. Threshold levels of the shown maps were individually adjusted to visualise flexible elements. Maps labelled with coloured boxes were used for structural model building. Map G and H derived from a merged dataset of apo and elongating human Pol III. Both RELION^57^ and CryoSPARC^60^ were used for data processing, as indicated.

**Extended Data Fig. 4.**
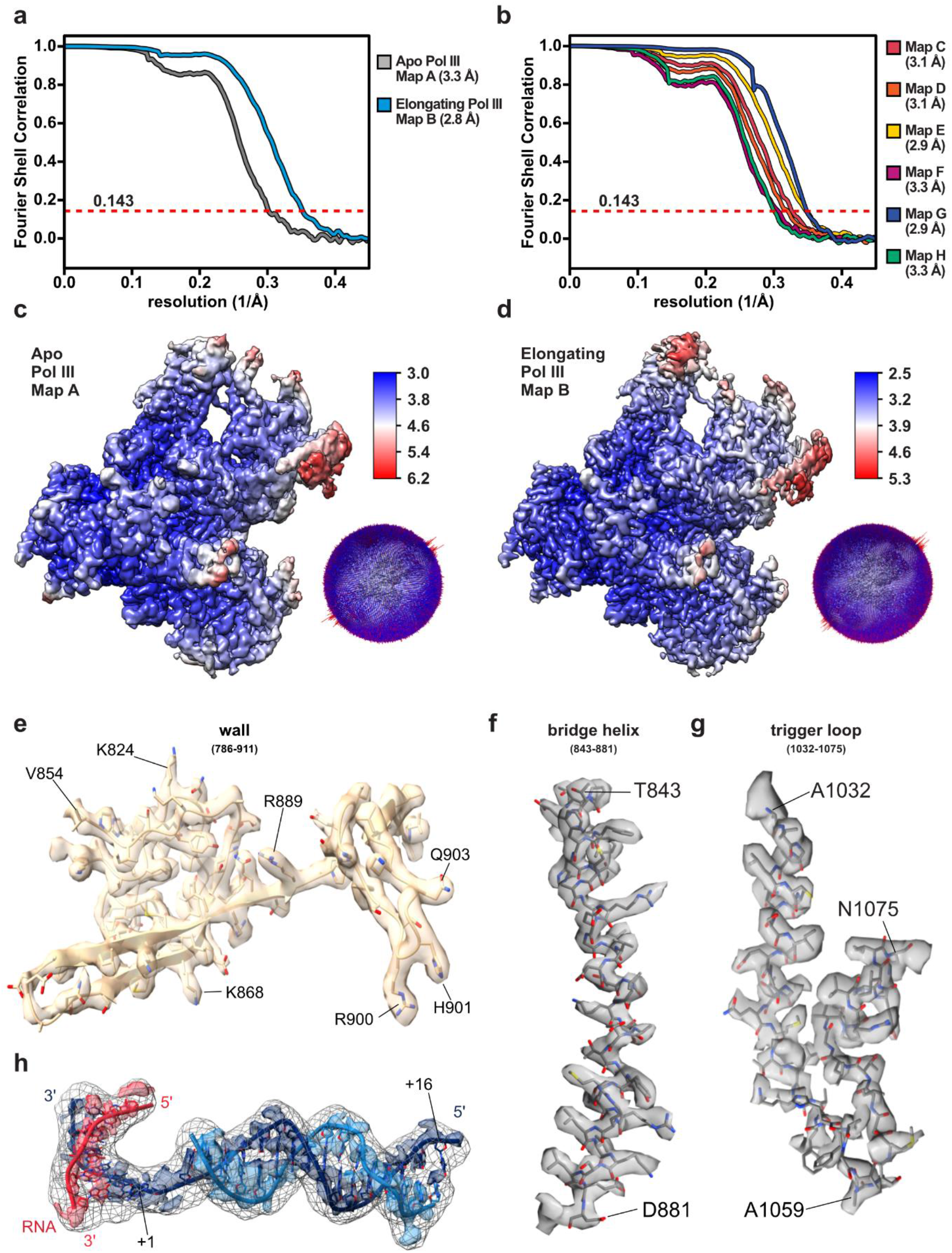
**a,** FSC curves of the apo and elongating Pol III showing final average resolutions of 3.3Å and 2.8Å, respectively (FSC = 0.143). **b**, FSC curves of cryo-EM maps with resolved flexible regions seen in Extended Data Fig. 3. Nominal resolution values for are given in brackets (FSC = 0.143). Local resolution estimation of **c**, apo Pol III and **d**, elongating Pol III as implemented in RELION. Corresponding angular distribution plots of all particles contributing to apo and elongating Pol III structures given on the right-hand side of **c**, and **d**, panels respectively. Fit of the modelled representative active site elements (stick representation): **e**, wall, **f**, bridge helix and **g**, trigger loop into its corresponding cryo-EM densities (surface representation). **h**, Fit of the nucleic acids into its corresponding cryo-EM density (coloured surface representation). Low-pass filtered by Gaussian filter = 1.5 density shown in mesh representation.

**Extended Data Fig. 5.**
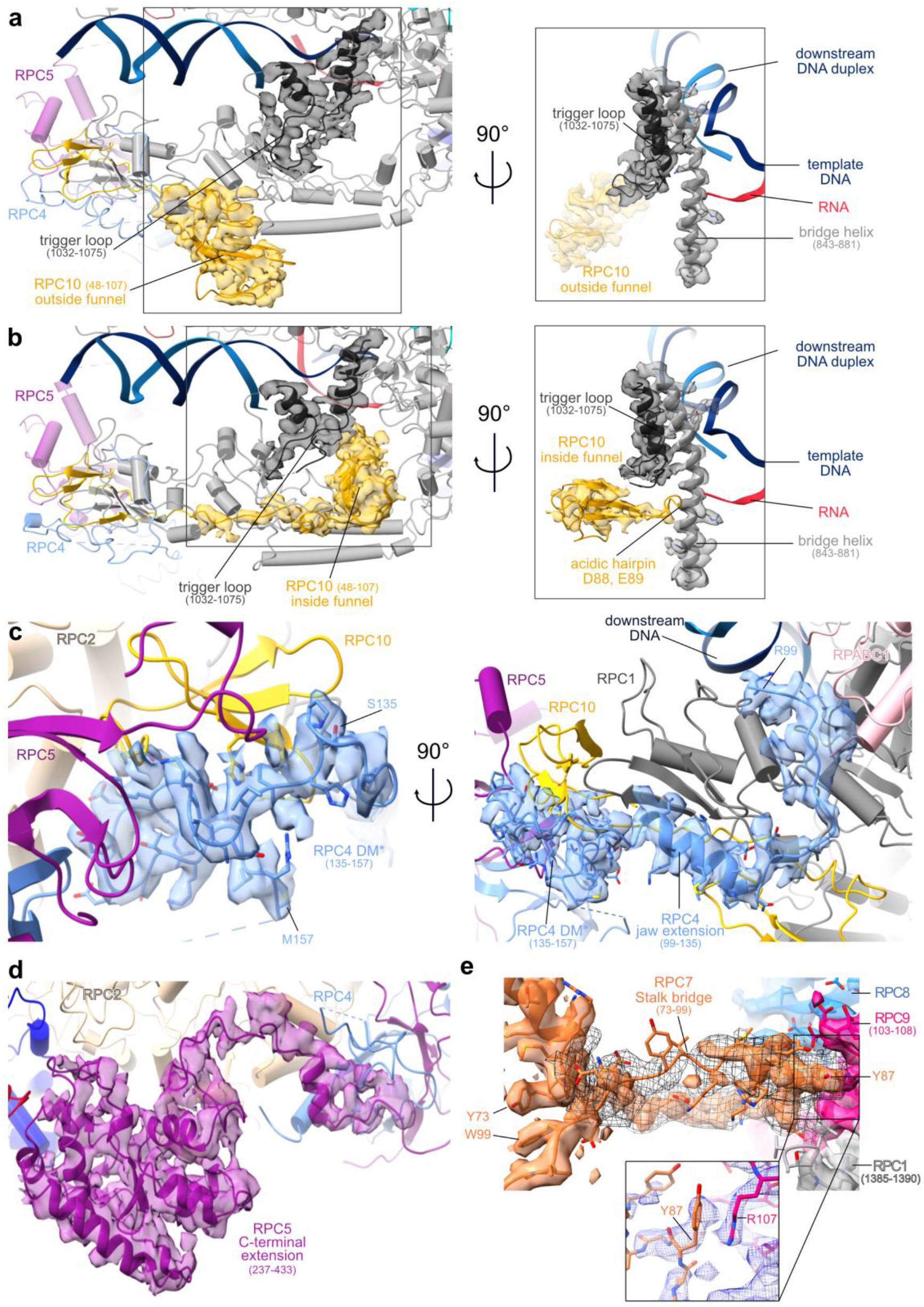
Cryo-EM density around newly built regions of human Pol III. **a,** RPC10 in the ‘outside funnel’ conformation. **b,** RPC10 in the ‘inside funnel’ conformation. On the left-hand side panel, surrounding environment is shown in transparent cartoon representation, and RPC2 and RPC1 bridge helix were removed for clarity. The right-hand side panel shows a 90° rotation of squared sections. **c,** Newly built RPC4 dimerization domain shown in molecular context (cartoon). **d,** RPC5 C-terminal domain, built into map F, is anchored to RPC2 and RPC4. **e,** Stalk bridge connecting the heterotrimer. Black mesh representation shows a 5 Å low-pass filtered cryo-EM density. Key interacting residues Y87 of RPC7 and R107 of RPC9 are shown in a close-up. The corresponding cryo-EM density is shown as blue mesh and belongs to map B2 (RELION multi-body refinement on elongating Pol III) sharpened by density modification.

**Extended Data Fig. 6.**
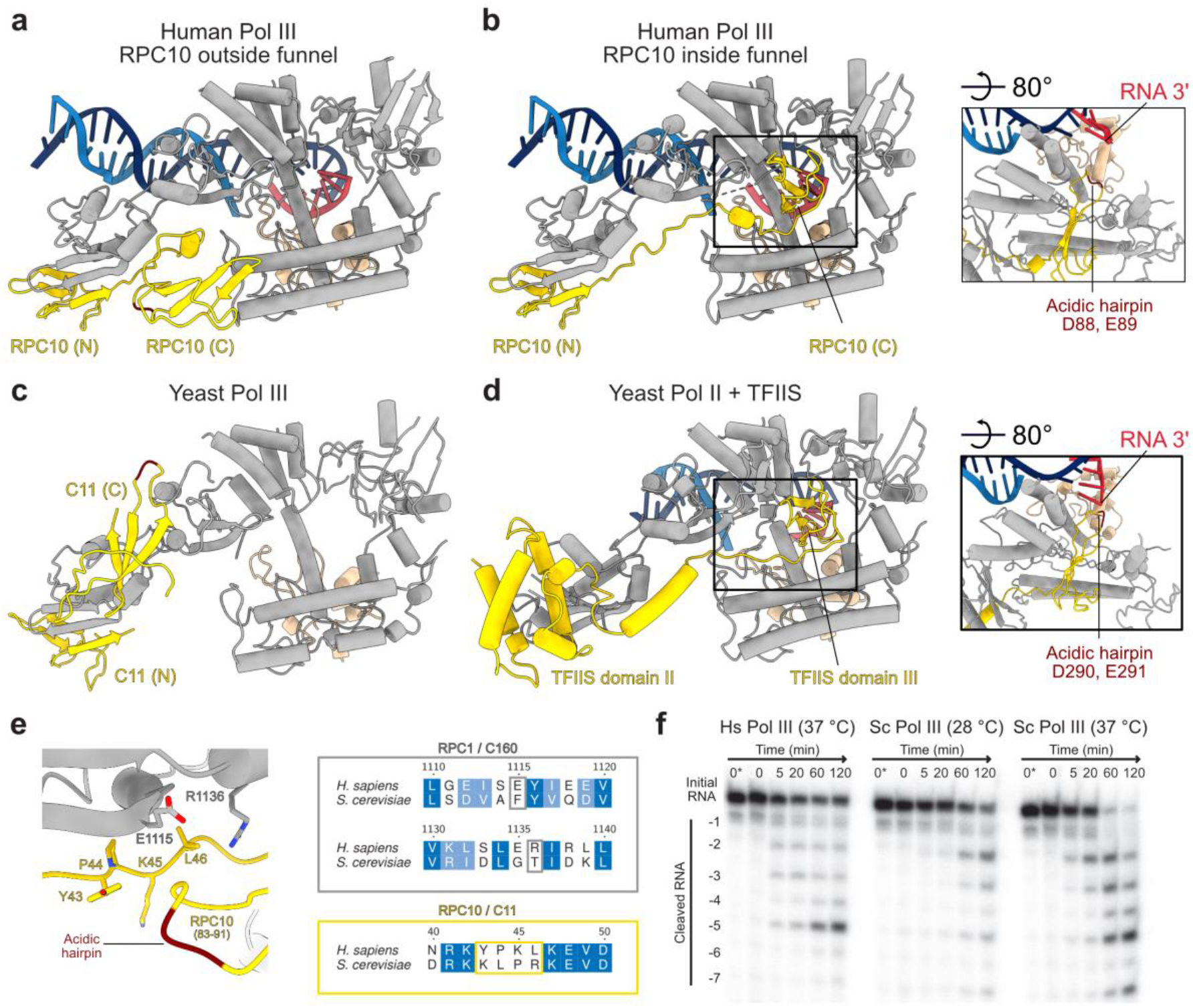
Structures of human RPC10 in different conformations. **a-d**, Shown are cartoon representations of human RPC10 or its homologues in yeast, of nucleic acids and of the polymerase regions that form the pore and funnel domains and the jaw domain. **a**, RPC10 C-ribbon (C) adopting the ‘outside funnel’ conformation in the elongating Pol III structure. **b**, RPC10 (C) adopting the ‘inside funnel’ conformation upon straightening of the linker. **c**, C11 (the yeast homolog to RPC10) shows a different conformation of its C-ribbon domain, whereas the position of the N-ribbon (N) is similar in both species. Shown is the yeast apo Pol III structure (PDB 5fja) because the C-ribbon is not visible other conformational states. **d**, Structure of yeast TFIIS bound to Pol II in its ‘reactivation intermediate’ conformation (PDB 3PO3). Acidic hairpins of RPC10, C11 and TFIIS are coloured dark red. Black frames in **a, d** highlight the position of the C-ribbons within the funnel and pore domain as shown by the rotated close-up views on the right. **e**, The RPC10 C-ribbon in the ‘outside funnel’ conformation folds back and bind its linker and the RPC1 jaw. RPC10 from residues 83 to 91 are shown as ribbon, and interacting residues within 4 Å distance are shown as sticks (left). Sequence alignments of *H. sapiens* RPC1 and *S. cerevisiae* C160 and of *H. sapiens* RPC10 and *S. cerevisiae* C11 covering the respective regions (right) shows that the contacting residues are not conserved. **f**, RNA cleavage assay of the ^32^P-labelled RNA annealed to the same transcription scaffold as used for structure determination of elongating Pol III. Shown is a time course from 0 to 120 minutes. Lane 0* contains RNA in the absence of polymerase. Hs – *H. sapiens*; Sc – *S. cerevisiae*. Image has been cropped for clarity.

**Extended Data Fig. 7.**
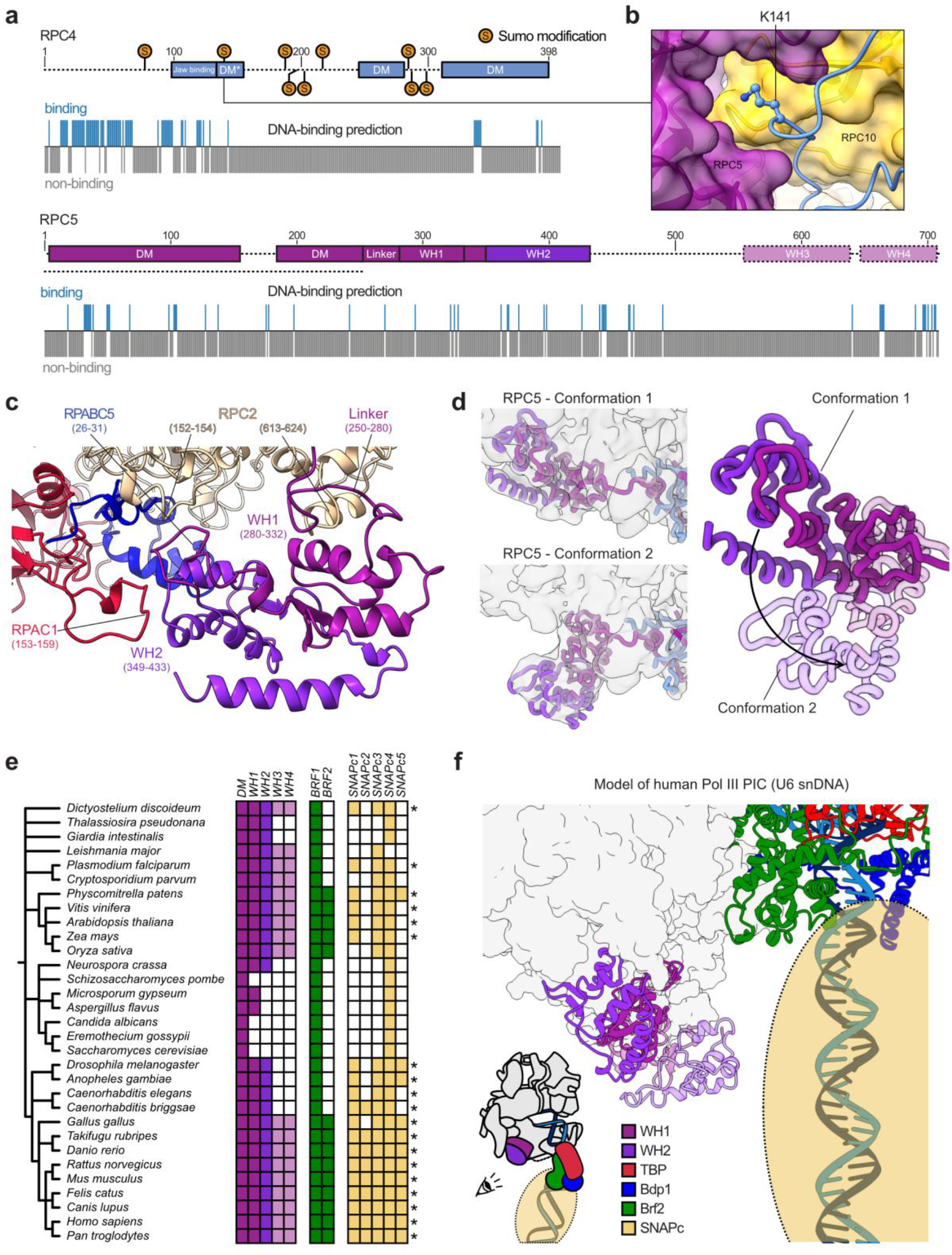
Structural and computational analysis of RPC4-RPC5 extensions. **a**, Schematic domain organisation of human RPC4 and RPC5 with DNA binding predictions below as predicted by DP-Bind^88^. Sumo-modifications of human RPC4 (K78, K141, K190, K199, K206, K220, K285, K288, K302) are marked on the sequence and were obtained from the PhosphoSitePlus database^89^. Dashed lines indicate regions that were not visible in the human Pol III cryo-EM maps. Human RPC5 contains, according to sequence homology prediction^90^, four WH domains that are absent in *S. cerevisiae*. **b**, Close-up view onto K141 of RPC4 being buried within the interface of RPC4 DM* (shown as cartoon), RPC5 and RPC10 (shown as transparent surfaces). **c**, Structural model of newly built RPC5 WH1 and WH2 domains binding the Pol III core subunits RPC2, RPAC1, and RPABC5. Corresponding contact points are labelled. **d**, Conformational dynamics of the WH1-WH2 domains. Left: Cryo-EM maps of human Pol III, in which RPC5 WH domains adopt conformation 1 (map F, top) and conformation 2 (map H, bottom). Cryo-EM maps were low-pass filtered to 6 Å and are shown as transparent surfaces. RPC4 and RPC5 are shown as thick ribbons. Right: Superimposition of the WH domains in the two conformations. The black arrow illustrates their movement. The WH domains in conformation 2 are shown as transparent cartoons **e,** Phylogenetic analysis of RPC5 and its WH domains, of TFIIIB components Brf1 and Brf2, as well as of the SNAPc genes. The colour code for RPC5 is the same as in **a**. Coloured and white squares indicate identified and not-identified homologues, respectively. The stars mark species, in which all the SNAPc building blocks were found that are likely required to form a functional SNAPc complex (SNAPc1, SNAPc3, SNAPc4) For further details of phylogenetic analysis see Methods. **f**, Model of the human Pol III PIC assembled onto the U6 snDNA promoter. Pol III core is shown as surface and RPC5 WH domains, Brf2, Bdp1, TBP and DNA are represented as cartoons. SNAPc is illustrated as transparent sphere. The model was generated using Chimera and Coot by: i) superimposing human TBP from PDB 5n9g onto yeast TBP in PDB 6f42, ii) extending the upstream DNA in 5n9g manually with ideal B-DNA carrying the human U6 snDNA sequence in Coot, iii) superimposing the elongating human Pol III model onto the model of yeast Pol III with Chain C (RPAC1) being used for alignment. SNAPc was placed onto the modelled downstream DNA according to information retrieved from the literature^33,91^.

**Extended Data Fig. 8.**
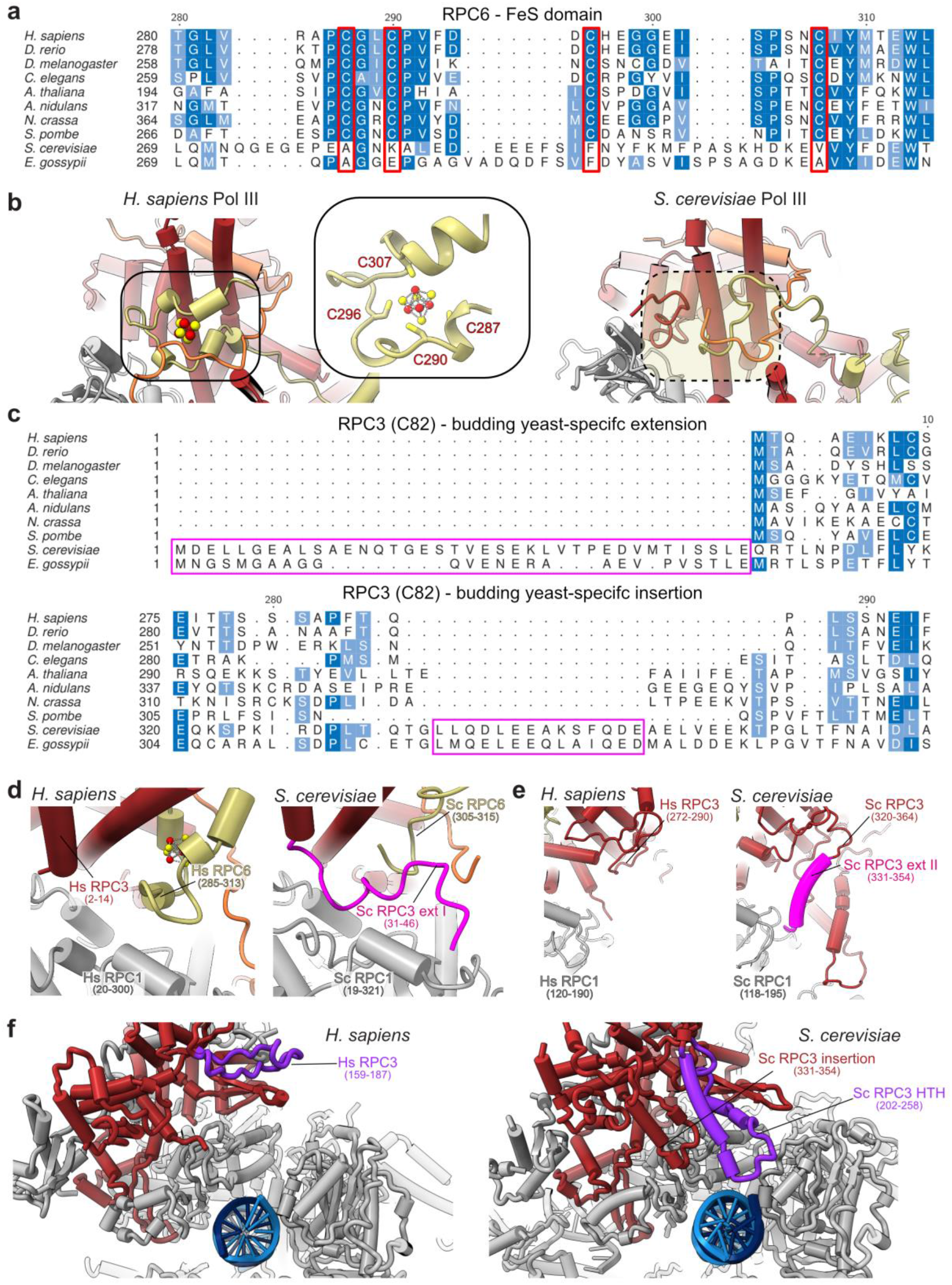
Structure, function and evolution the RPC6 FeS domain and RPC3 extensions. **a**, Multiple sequence alignment (MSA) of the RPC6 C-terminal FeS domain from model organisms. MSA was generated via T-Coffee^92^ implemented into the EMBL-EBI bioinformatic tools web service^93^. The four cysteine residues required for coordination of the FeS cluster are highlighted in red. **b**, Comparison of the RPC3-RPC6-RPC7 heterotrimer from *H. sapiens* (left, elongating Pol III) and *S. cerevisiae* (right, PDB 6eu3). The cubane FeS cluster is coloured in red and yellow. The position of the FeS cluster is framed, and a close-up view shows the four cysteines coordinating the FeS cluster. The equivalent position in *S. cerevisiae* Pol III structure, which lacks the FeS cluster, is marked with a dashed oval. **c**, MSA of the RPC3 (C82 in *S. cerevisiae*) N-terminus and of region 275-294 (*H. sapiens*) generated with T-Coffee. The MSA of the RPC3 N-terminus was manually adjusted, based on the available structural information (see panel **d**), so that the N-terminus of budding yeasts becomes apparent as an N-terminal extension instead of an insertion. The budding yeast specific N-terminal extension and central insertion (331-354 in *S. cerevisiae*) are highlighted with pink frames. **d-f**, Close-up comparisons between *H. sapiens* (elongating Pol III) and *S. cerevisiae* (PDB 6eu3 in **d,e** and 6f41 in **f**) Pol III structural elements that contribute to heterotrimer integrity and anchoring to the RPC1 (C160 in *S. cerevisiae*) core subunit. **d**, Budding yeast-specific N-terminal extension is positioned similarly as the FeS cluster in *H. sapiens* Pol III. Notably, only 16 residues (31-46) of the extension are visible, whereas the remaining amino acids are flexible. **e**, The central insertion (331-354) binds the polymerase core, whereas this contact point is missing in *H. sapiens*. **f**, *S. cerevisiae* Pol III contains a helix-turn-helix (HTH) insertion (202-258) that binds the downstream DNA in the *S. cerevisiae* PIC (PDB 6f41). The HTH insertion further binds and stabilises the central insertion (331-354) and, thereby, likely contributes to the anchoring of the heterotrimer to the polymerase core. Both elements are absent in *H. sapiens* although the counterpart region (159-187 in *H. sapiens*) has a similar shape (albeit without α-helical elements) that points into a different direction. Hs – *H. sapiens*; Sc – *S. cerevisiae*. Sc RPC1, Sc RPC3, Sc RPC6 relate to Sc C160, Sc C82, Sc C34, respectively, according to the commonly used nomenclature of *S. cerevisiae* Pol III subunits.

**Extended Data Fig. 9.**
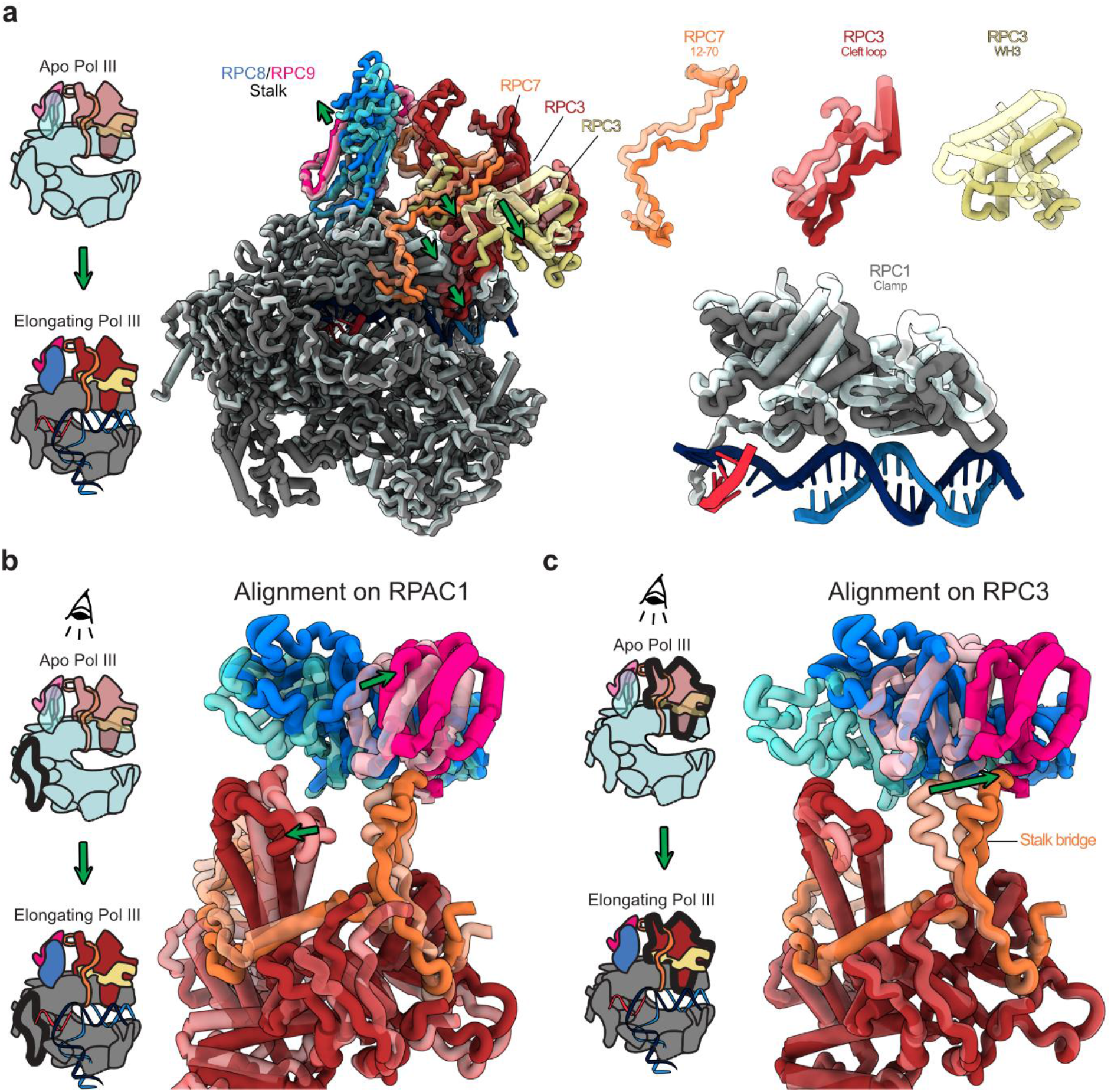
RPC7 tethers heterotrimer and stalk that move in opposite directions. Protein elements are shown as thick ribbon to better visualize conformational changes. **a**, Left: Schematic showing the transition from DNA-unbound Apo Pol III to DNA/RNA-bound elongating Pol III. Middle: Side view showing the superimposition of apo and elongating human Pol III with RPAC1 being used for aligning the two structures. Apo Pol and elongating human Pol III are shown as transparent and solid ribbons, respectively. Green arrows illustrate movements of Pol III heterotrimer and clamp and the polymerase stalk domain. Right: Close-up views on selected moving elements showing that heterotrimer and clamp move towards the bound downstream DNA upon transition from apo to elongating Pol III. **b, c**, Top views onto the heterotrimer and stalk domains with two different subunits (RPAC1 in **b** and RPC3 in **c**) used to align apo and elongating human Pol III structures. Aligned subunits are outlined in bold in the associated schematics shown on the left. **b**, The RPC7 stalk bridge tethers heterotrimer and stalk although they the two domains move in opposite directions. **c**, The tip of the RPC7 stalk bridge is anchored to the stalk domain and moves in the same direction.

**Extended Data Fig. 10.**
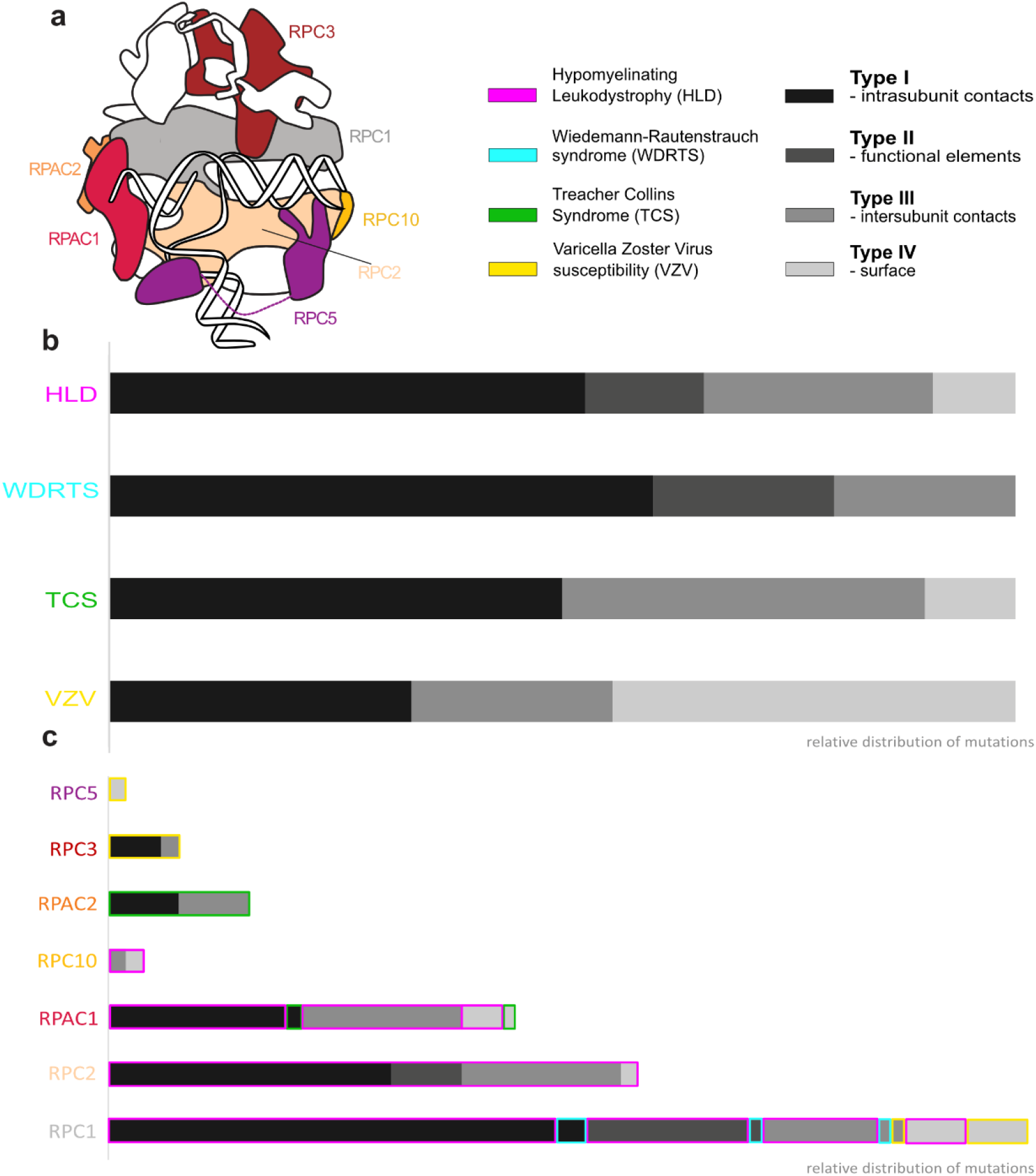
Outlined types of point mutations in Pol III correlated with diseases. **a,** Cartoon representation of Pol III showing affected subunits (in colour). Diseases correlated with mutations and types they have been assigned to. **b,** Relative distribution of types of mutations in each considered disease. **c,** Relative distribution of types of mutations in each subunit. Coloured outlines reference diseases. Size of bars corresponds to the number of mutations within each subunit. For the type assignment, see Methods. For details of individual mutations, see Supplementary Table 2.

