## Supplementary Information Figures and Tables for "Cryo-EM structures of human RNA polymerase III in its unbound and transcribing states"

***H. sapiens* (Hs) RPC7α - *S. cerevisiae* RPC7/C31**

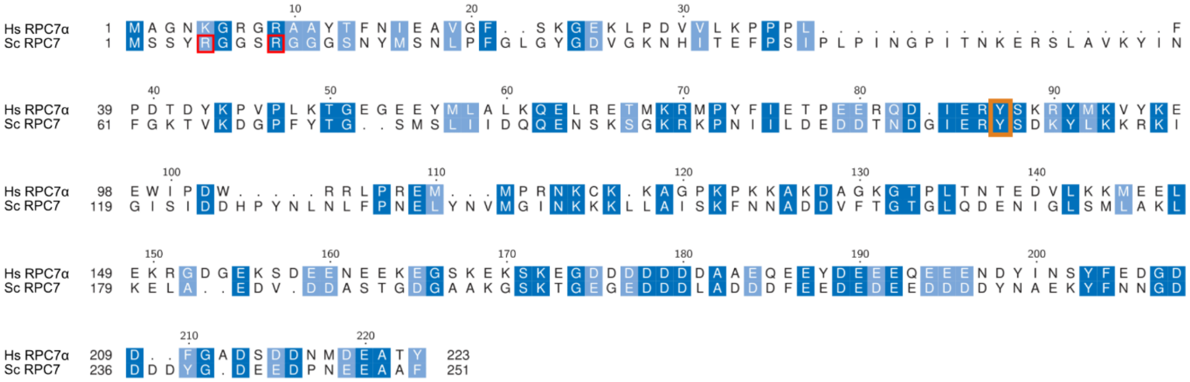

| Range (Hs) | Sequence identity (%) |
|---|---|
| 1-25 | 20 |
| 1-223 | 29 |

Reported methylation sites

Anchor tyrosine

***H. sapiens* (Hs) RPC7β - *S. cerevisiae* (Sc) RPC7/C31**

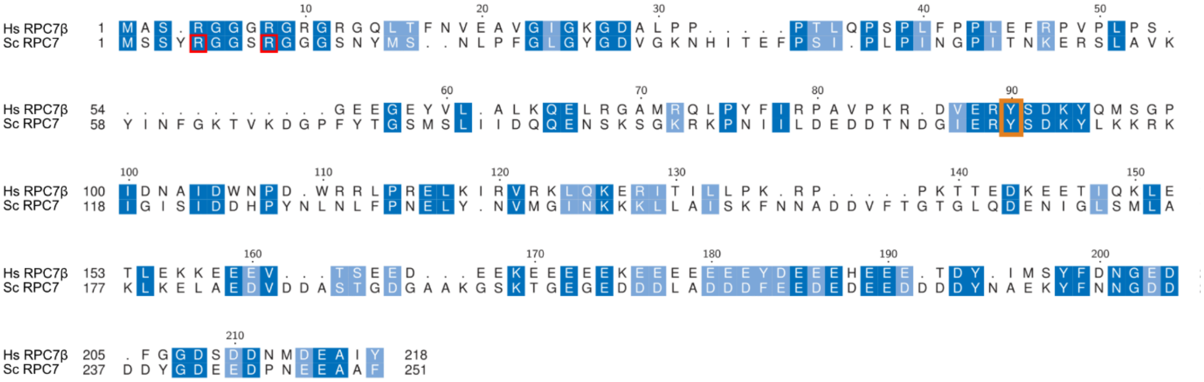

| Range (Hs) | Sequence identity (%) |
|---|---|
| 1-25 | 43 |
| 1-223 | 29 |

Reported methylation sites

Anchor tyrosine

**Supplementary Figure 1 | Sequence alignment for *H. sapiens* RPC7α/β to *S. cerevisiae* RPC7/C31.** Top: *H. sapiens* RPC7α aligned to *S. cerevisiae* RPC7/C31. Bottom: *H. sapiens* RPC7α aligned to *S. cerevisiae* RPC7/C31. Sequence identities of the protein N-termini (1-25) and of the full-length proteins are shown below the sequence alignments. Red frames indicate reported[1] arginine methylation sites in *S. cerevisiae* RPC7/C31. Orange frames highlight the anchoring tyrosine (F87 in RPC7α), which connects Pol III heterotrimer and stalk (main text).

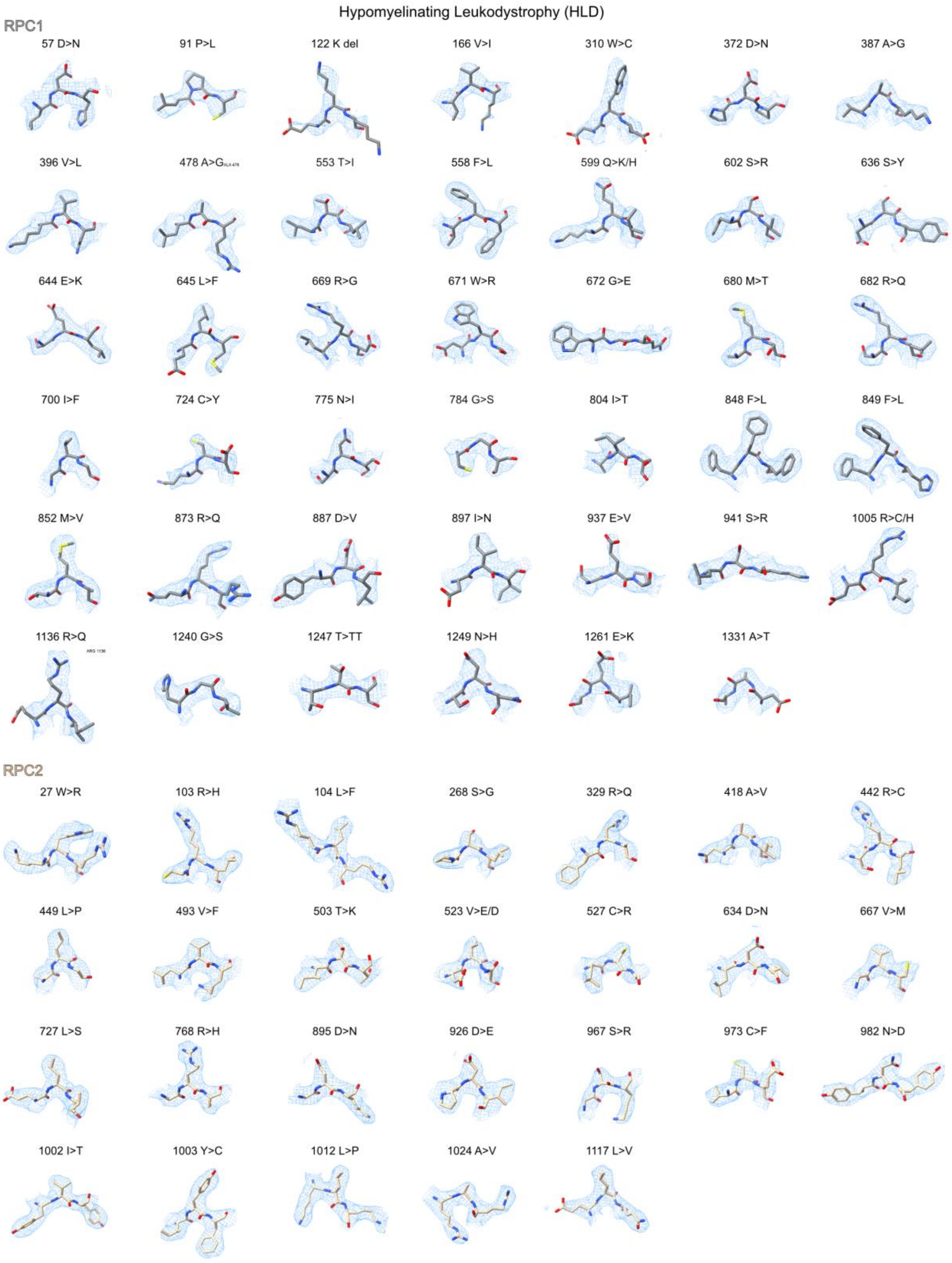

**Supplementary Figure 2 | Cryo-EM density fits of Pol III disease-associated mutations.** Shown are residues, mutated in genetic diseases (see **Supplementary Table 2)**, of the human Pol III model (EC-1 Pol III) and their fit into cryo-EM density (map B). Residues are displayed with flanking residues (-1,1) and the same threshold level (0.04) of map B was used for all shown fits.

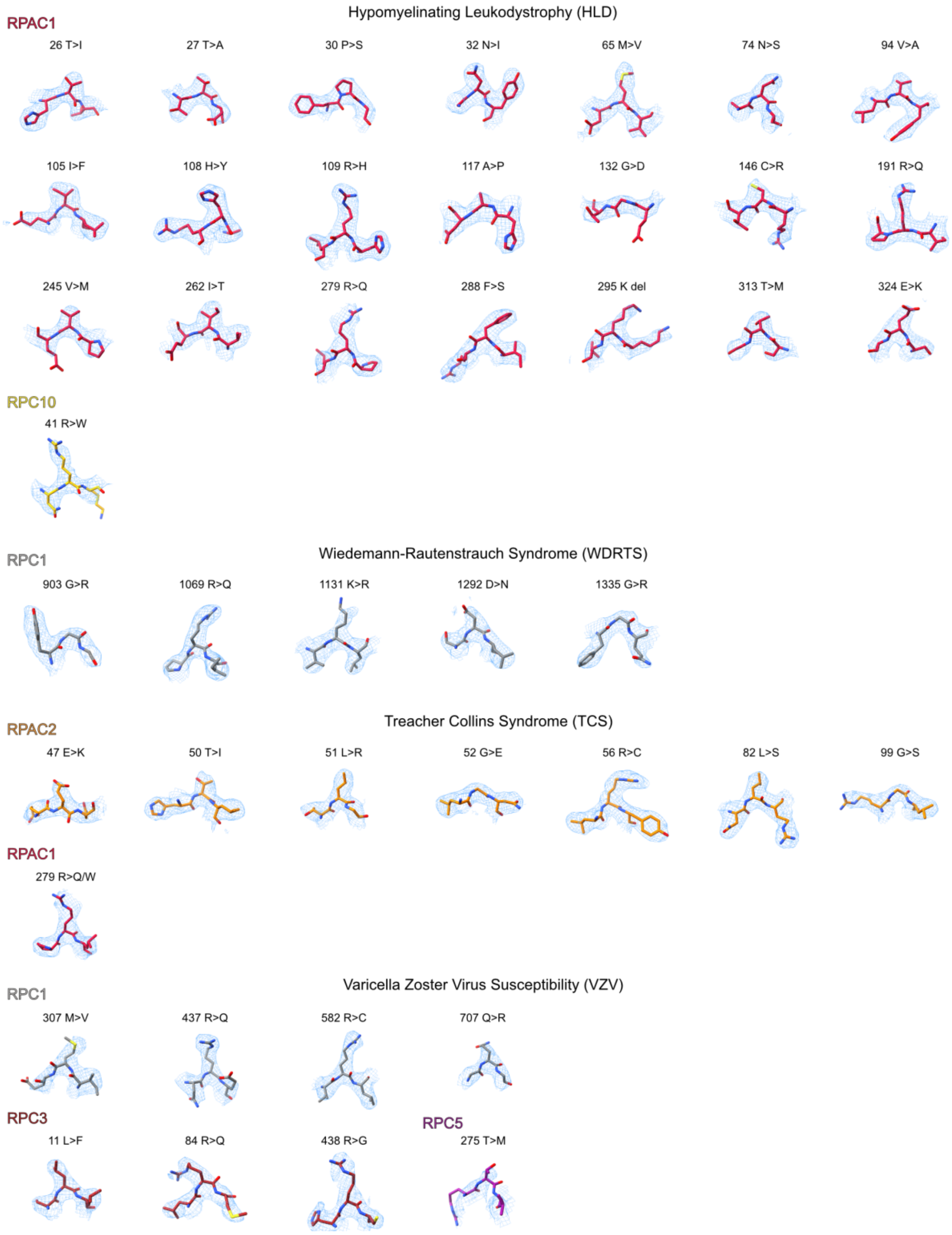

**Supplementary Figure 2 (continued)**

**Supplementary Table 1 | Phylogenomic analysis of Pol III RPC5 subunits and Pol III basal transcription factors.** Listed are the analysed species, proteome identifiers and UniProt identifiers for homologs of RPC5 (Pol III), BRF1 and BRF2 (basal transcription factor TFIIIB) and SNAPc1, SNAPc2, SNAPc3, SNAPc4, SNAPc5 (basal transcription factor SNAPc). Homologs were retrieved using the HMMER[1] tool. See Extended Data Fig. 7e for the corresponding phylogenomic tree.

| Species | Proteome ID | RPC5 | BRF1 | BRF2 | SNAPc1 | SNAPc2 | SNAPc3 | SNAPc4 | SNAPc5 |
|---|---|---|---|---|---|---|---|---|---|
| *Dictyostelium discoideum* | UP000002195 | Q86JI9 | Q54U85 | | Q54B89 | | Q555K3 | Q54NA6 | |
| *Thalassiosira pseudonana* | UP000001449 | B8BWK0 | B8BTG5 | | | | | B8CFV0 | |
| *Giardia intestinalis* | UP000001548 | A8BX96 | E2RTV0 | | | | | E2RTS5 | |
| *Leishmania major* | UP000000542 | Q4QG17 | Q4QA51 | | | | E9AFU3 | | |
| *Plasmodium falciparum* | UP000001450 | Q8IKP3 | A0A144A465 | | Q8I600 | | Q8IKM4 | Q7KQL1 | |
| *Cryptosporidium parvum* | UP000006726 | Q5CW66 | Q5CRA9 | | | | A3FPM6 | Q5CUX4 | |
| *Physcomitrella patens* | UP000006727 | A9RV70 | A0A2K1K844 | A0A2K1JX89 | A0A2K1J6Z8 | | A0A0D6CBM7 | A0A2K1JLB9 | A0A2K1IMI5 |
| *Vitis vinifera* | UP000009183 | D7SPH0 | D7TL42 | D7TJK6 | D7TC57 | | D7T5Z1 | D7U7H5 | |
| *Arabidopsis thaliana* | UP000006548 | Q9FGZ3 | F4IW19 | O81787 | Q8RXK3 | | Q8L627 | Q9LV31 | |
| *Zea mays* | UP000007305 | UPI0002207C81 | A0A1D6MBH4 | A0A1D6PYC9 | B4FT72 | | A0A1D6G6D0 | A0A1D6HS21 | |
| *Oryza sativa* | UP000007015 | A2YT86 | B8AWC3 | A2YXG0 | | | A2WY80 | A2YI13 | |
| *Neurospora crassa* | UP000001805 | Q7S9I7 | Q7RY40 | | | | | Q1K6E3 | |
| *Schizosaccharomyces pombe* | UP000002485 | O74883 | Q9P6R0 | | | | | P39964 | |
| *Microsporum gypseum* | UP000002669 | E4UVQ2 | E5R1V4 | | | | | E4V5A4 | |
| *Aspergillus flavus* | UP000001875 | B8N1U9 | B8N559 | | | | | B8MY80 | |
| *Candida albicans* | UP000000559 | A0A1D8PQS2 | P43072 | | | | | Q5A683 | |
| *Eremothecium gossypii* | UP000000591 | Q750I4 | Q758H5 | | | | | Q753L5 | |
| *Saccharomyces cerevisiae* | UP000002311 | P36121 | P29056 | | | | | P22035 | |
| *Drosophila melanogaster* | UP000000803 | Q9VP85 | Q9VEL2 | | Q9VF25 | | Q7JUY8 | Q9VHX0 | Q8INC7 |
| *Anopheles gambiae* | UP000007062 | Q7Q6C6 | Q7QB20 | | A7UUY9 | | Q7QFT6 | A7UVR7 | F5HKF9 |
| *Caenorhabditis elegans* | UP000001940 | Q23206 | A0A131MCT5 | | Q7JNN6 | | O44168 | P91868 | |
| *Caenorhabditis briggsae* | UP000008549 | A8XLK1 | A8XH51 | | A8XT63 | A8WWB8 | A8XX82 | A8XFC6 | |
| *Gallus gallus* | UP000000539 | E1C9J0 | F1NQ85 | E1C4I7 | F1NJK3 | | F1NW98 | A0A1L1S0G0 | Q5ZI06 |

### Supplementary Table 1 (continued)

| Species | Proteome ID | RPC5 | BRF1 | BRF2 | SNAPc1 | SNAPc2 | SNAPc3 | SNAPc4 | SNAPc5 |
|---|---|---|---|---|---|---|---|---|---|
| *Takifugu rubripes* | UP000005226 | A0A3B5KDH1 | H2UWY0 | A0A3B5JWS9 | H2S607 | H2SIN9 | H2TJN2 | A0A3B5KEC1 | A0A3B5KKF9 |
| *Danio rerio* | UP000000437 | F1R866 | Q5TZ89 | A8KBY2 | Q7ZUP7 | E7F685 | F1QWM0 | F1QVS5 | Q1LXI0 |
| *Rattus norvegicus* | UP000002494 | D3ZP45 | D4A8W8 | Q4V8D6 | F7ES94 | Q68FX5 | Q5BK68 | D4A3C9 | D3ZTK9 |
| *Mus musculus* | UP000000589 | Q9CZT4 | Q8CFK2 | Q3UAW9 | Q8K0S9 | Q91XA5 | Q9D2C9 | Q8BP86 | Q8R2K7 |
| *Felis catus* | UP000011712 | A0A2I2V2B4 | A0A337S677 | M3X9N0 | M3W8U2 | M3WLN3 | M3VZV1 | A0A5F5XJV4 | M3WGN7 |
| *Canis lupus* | UP000002254 | E2RHN7 | F1PRK4 | E2R245 | E2QXT2 | J9NTT0 | E2REJ9 | J9NSI5 | E2RR56 |
| *Homo sapiens* | UP000005640 | Q9NVU0 | Q92994 | Q9HAW0 | Q16533 | Q13487 | Q92966 | Q5SXM2 | O75971 |
| *Pan troglodytes* | UP000002277 | K7BW50 | H2Q908 | H2QW11 | H2Q8F4 | H2RHY8 | H2R1P5 | H2QY63 | H2Q9N3 |

**Supplementary Table 2 | Disease causing point mutations in Pol III.** In bold mutations have been reported as homozygous. SASA – Solvent Accessible Surface Area. Average for whole complex: SASA - 42.4 Å$^2$; degree - 15.

| Related disease | Subunit | Mutation | Side chain contacts | SASA (Å$^2$) | Residue degree | Type |
|---|---|---|---|---|---|---|
| Hypomyelinating Leukodystrophy (HLD) | RPC1 | 57 D>N[3] | RPC1: I34, M60, L52, G54, H58 R59 | 7.3 | 19 | I |
| | | 91 P>L[4] | RPC1: P220, L221, L224, I251 | 9.5 | 14 | I |
| | | 122 K deletion[5] | RPC1: I115, L117, S118, L237 | 40.4 | 20 | I |
| | | **166 V>I**[4] | RPC1: L105, Q106, K110, I174, H176 | 7.7 | 16 | II |
| | | 310 W>C[6] | RPC1: L90, P91, P220, L221, I306, M307 | 12.1 | 24 | I |
| | | 372 D>N[7] | RPC1: N374, L375, R487 RPAC2: H74 | 0.9 | 20 | III |
| | | 387 A>G[4] | RPC1: P383, H415, L454, V480 | 0.0 | 21 | I |
| | | 396 V>L[8] | RPC1: I401, L404, K445, D448, V450 | 0.6 | 18 | I |
| | | 478 A>G[9] | RPC1: V382, L454 | 0.1 | 15 | I |
| | | 553 T>I[5] | RPC1: A549, F601, S649 | 0.0 | 20 | I |
| | | 558 F>L[7] | RPC1: D556, T557, I588, L594, T596, RPABC3: R24, N44, L121 | 27.1 | 16 | III |
| | | 599 Q>K/**H**[5] | RPC1: T587, W595, T596 | 26.5 | 19 | I |
| | | 602 S>R[4,10] | RPC1: Q599, S643, E644 | 13.8 | 18 | I |
| | | 636 S>Y[7] | RPC1: G621, Q623, Y624, N634 | 0.4 | 17 | I |
| | | 644 E>K[5] | RPC1: Q641, S643 RPABC3: R140 | 34.1 | 18 | II |
| | | 645 L>F[5] | RPC1: T553, K598, F601, T639, I640L, S647, G648, M650 | 6.1 | 17 | I |
| | | 669 R>G[11] | RPC1: V530, Y665, A910, A911, E913 | 55.0 | 17 | I and IV |
| | | **671 W>R**[4] | RPC1: R614, L667, D670, E675 | 79.0 | 15 | I and IV |
| | | **672 G>E**[5,7] | - | 28.4 | 11 | I |
| | | 680 M>T[5] | RPC1: I377, L529, I539, A540, F664, A676 | 1.9 | 22 | I |
| | | 682 R>Q[9] | RPC1: D576, I604, P607, D678 | 16.6 | 21 | I |
| | | 700 I>F[3] | RPC1: 703, RPC2: V940, I944, L947, L984, Y1003 | 1.1 | 18 | II |
| | | **724 C>Y**[7] | RPC1: Y721, E755 | 0.3 | 17 | II |
| | | 775 N>I[7] | RPC1: D702, L771, D772, P777, L778 | 11.5 | 20 | I |
| | | 784 G>S[10] | - | 15.1 | 10 | I |
| | | 804 I>T[6] | RPC1: A803,V809, M852, A853 | 19.7 | 12 | II |
| | | **848 F>L**[5] | RPC1: T845, RPC2: P473, W476, P481, L664 | 0.0 | 21 | II |
| | | **849 F>L**[5] | RPC1: V809, S817, L818, G832, T845, M852 | 1.2 | 20 | II |
| | | 852 M>V[4,7,12] | RPC1: V809, F848, F849 RPC2: L471, P473 | 9.9 | 22 | II |
| | | 873 R>Q[11] | RPC1: R353, G869, Y870, F1321 | 18.2 | 19 | II |
| | | 887 D>V[5] | RPC1: Q885, T889, R891, RPABC2: P111 | 33.2 | 14 | III |

| | | | | | | |
|---|---|---|---|---|---|---|
| Hypomyelinating Leukodystrophy (HLD) | RPC1 | 897 I>N[13] | RPC1: I898, Q899, F900 RPABC1: L165 | 0.4 | 17 | III |
| | | 937 E>V[5] | RPC1: C934, T1007, T1009 | 49.6 | 16 | IV |
| | | 941 S>R[8] | RPC1: D991, E944 | 9.1 | 18 | I |
| | | 1005 R>C/H[4,7,13,14] | RPC1: C934, P935, A939, Y1001, T1007 | 38.8 | 16 | I |
| | | 1136 R>Q[8] | RPC1: S1133, RPC10: L46, K47 | 1.9 | 20 | III |
| | | 1240 G>S[5] | - | 6.4 | 9 | I |
| | | 1247 T>TT[7] | RPC1: T1246 | 44.3 | 12 | I and IV |
| | | 1249 N>H[12] | RPC1: R1069, I1084, E1270 | 31.3 | 14 | I |
| | | 1261 E>K[4] | RPABC1: P146, I193, R195, R207 | 6.2 | 17 | III |
| | | 1331 A>T[14] | RPC1: F20, V1315, H1327 | 0.0 | 17 | I |
| | RPC2 | **27 W>R**[5] | RPC2: R28, P31, E660 | 51.4 | 16 | I and IV |
| | | 103 R>H[5] | RPC2: H100, L140, E160, Y168, I170, E175, R704 | 12.6 | 23 | I |
| | | 104 L>F[4] | RPC2: H100, I705, RPABC4: I43, Y45 | 2.6 | 20 | II |
| | | 268 S>G[4] | RPC2: V237, F265 | 1.5 | 15 | I |
| | | 329 R>Q[5] | RPC2: N327, N524, L526, C527, RPC5: R136 | 26.5 | 20 | III |
| | | **418 A>V**[5] | RPC2: L372, G414 | 7.0 | 17 | I |
| | | 442 R>C[4] | RPC2: D164, S441, I697, R704 | 29.8 | 15 | II |
| | | 449 L>P[5] | RPC2: V40, V667, I728 | 10.4 | 17 | I |
| | | 493 V>F[5] | RPC2: R454, P485 | 25.0 | 15 | I |
| | | 503 T>K[15] | RPC2: F571, G589, R591, C593 | 1.5 | 17 | II |
| | | **523 V>E/D**[4,5,15] | RPC2: I333, L526 | 10.9 | 12 | I |
| | | 527 C>R[4] | RPC2: E250, Q253, E529, RPC4: F356, RPC5: P31 | 0.7 | 17 | III |
| | | 634 D>N[9] | RPC2: S573, R591, L592, V635 | 2.7 | 16 | I |
| | | 667 V>M[5] | RPC2: L39, L498 | 3.8 | 11 | I |
| | | 727 L>S[5] | RPC2: L30, L34, L498, K723 | 38.6 | 17 | I |
| | | 768 R>H[13] | RPC2: A713, G765, C769, T888, R889, R890 RPAC1: H108 | 5.9 | 20 | III |
| | | 895 D>N[5] | RPC2: P891, E892, M1015 | 4.4 | 19 | I |
| | | 926 D>E[13] | RPC2: T740, S761, RPABC5: T9 | 0.3 | 21 | III |
| | | 967 S>R[5] | RPC2: Y674, H959 | 23.2 | 10 | I |
| | | 973 C>F[5] | RPC2: V969, Y983 | 8.1 | 20 | I |
| | | 982 N>D[5] | RPC2: Y983, K986, RPAC1: S284, R285 | 14.7 | 15 | III |
| | | **1002 I>T**[5] | RPC2: A742, M744, V989, Y1001 RPC1: S697 | 0.0 | 21 | I and III |
| | | **1003 Y>C**[5] | RPC2: L943, L946, L947, L984, RPC1: S697, I698, G699, I700 | 0.0 | 20 | I and III |
| | | 1012 L>P[5] | RPC2: M689, K896, S898 | 0.2 | 17 | II |
| | | 1024 A>V[5] | RPC1: F363, I389 | 22.1 | 13 | III |
| | | 1117 L>V[5] | RPC2: L1113, M1120, L1122, RPC1: L345, L1328 | 0.4 | 21 | I and III |
| | RPAC1 | **26 T>I**[16,17] | RPAC1: D28, D69, RPC2: E996 | 9.1 | 14 | I and III |
| | | 27 T>A[17] | RPAC1: Y33, RPC1: D560, | 15.8 | 16 | I |

| | | | | | | |
|---|---|---|---|---|---|---|
| Hypomyelinating Leukodystrophy (HLD) | RPAC1 | 30 P >S[17] | RPAC2: M60, K61 | 6.4 | 13 | III |
| | | **32 N>I**[16,17] | RPAC1: T27, F29, Y33<br>RPC1: D560 | 3.1 | 18 | III |
| | | 65 M>V[16,17] | RPAC1: F49, F63, I68, F76,<br>RPAC2: V110 | 3.9 | 18 | III |
| | | **74 N>S**[16,17] | RPAC1: A70, R77, R78,<br>RPC2: G995 | 3.6 | 22 | III |
| | | 94 V>A[16,17] | RPAC1: V92, N97, L210,<br>RPABC4: R48, F55 | 2.3 | 16 | III |
| | | 105 I>F[17] | RPAC1: Q102, RPC2: A713 | 3.2 | 17 | III |
| | | 108 H>Y[17] | RPAC1: E104, I105, L112,<br>RPC2: E763, R764, G765 | 3.6 | 21 | III |
| | | 109 R>H[16,17] | RPAC1: I105, L198, I199<br>RPC2: Y714,<br>RPABC5: I2, I3, P4, V5 | 1.3 | 23 | III |
| | | 117 A>P[17] | RPAC1: M87, I115, L138, I190 | 0.3 | 17 | I |
| | | 132 G>D[16,17] | - | 2.6 | 8 | I |
| | | 146 C>R[16,17] | RPAC1: H166, L202, R203, P204 | 0.4 | 17 | I |
| | | 191 R>Q[16,17] | RPAC1: H116, G188, P192 | 101.9 | 14 | IV |
| | | 245 V>M[16,17] | RPAC1: E243, A249, A250, L253,<br>V296 | 10.0 | 17 | I |
| | | 262 I>T[16,17] | RPAC1: L253, S254, F257, A273,<br>V275 | 17.8 | 16 | I |
| | | **279 R>Q**[17] | RPAC1: V54, D64, N277 | 65.3 | 14 | I and IV |
| | | **288 F>S**[18] | RPAC1: E18, K294 | 37.2 | 14 | I |
| | | 295 K deletion[16,17] | - | 144.4 | 8 | IV |
| | | 313 T>M[17] | RPAC1: P227, V228, A229 | 7.5 | 13 | I |
| | | 324 E>K[16,17] | RPAC1: R120, V320 | 36.5 | 18 | I |
| | RPC10 | **41 R>W**[19] | RPC10: Y43, RPC1: Y1116,<br>E1118 | 91.4 | 12 | III and IV |
| Wiedemann-Rautenstrauch syndrome (WDRTS) | RPC1 | 903 G>R[20] | - | 1.1 | 8 | I |
| | | 1069 R>Q[20] | RPC1: T1064, L1065, G1066,<br>E1072, I1080, N1249, E1270,<br>T1274 | 12.6 | 24 | II |
| | | 1131 K>R[20] | RPC1: E1115, Y1116, V1172,<br>RPC4: K119 | 17.7 | 17 | III |
| | | 1292 D>N[20] | RPC1: I898, Q899, R1264, M1288 | 4.7 | 22 | I |
| | | 1335 G>R[20] | - | 2.1 | 11 | I |
| Treacher Collins Syndrome (TCS) | RPAC1 | 279 R>Q/W[16,17,21] | RPAC1: V54, D64, N277 | 65.3 | 14 | I and IV |
| | RPAC2 | 47 E>K[21] | RPAC2: H45, E46, L51 | 3.4 | 15 | III |
| | | 50 T>I[21] | RPAC2: D48,<br>RPAC1: R78, I79, K332 | 2.0 | 16 | III |
| | | 51 L>R[21] | RPAC2: E47, RPAC1: K329, F336 | 0.0 | 16 | III |
| | | 52 G>E[21] | - | 0.1 | 13 | I |
| | | 56 R>C[21] | RPAC2: G52, C68, G69,<br>RPC1: A562, Q566 | 19.9 | 22 | I and III |
| | | 82 L>S[21] | RPAC2: K55, I59, C68, Y70, F96, | 0.1 | 20 | I |
| | | 99 G>S[21] | - | 1.2 | 14 | I |

| | | | | | | |
|---|---|---|---|---|---|---|
| Varicella Zoster Virus Susceptibility (VZV) | RPC1 | 307 M>V[22] | RPC1: T303, W310, RPC3: L406, Y422 | 18.5 | 18 | III |
| | | 437 R>Q[22] | V408, Q409, G411, L432 | 68.2 | 19 | IV |
| | | 582 R>C[23] | RPC1: D609 | 89.2 | 12 | IV |
| | | 707 Q>R[22] | - | 126.8 | 9 | IV |
| | RPC3 | 11 L>F[22] | RPC3: K7, I447 | 10.5 | 20 | I |
| | | 84 R>Q[22] | RPC3: S80, R87, H243 | 20.7 | 18 | I |
| | | 438 R>G[22] | RPC3: R357, L435, RPC6: F293 | 19.6 | 18 | I and III |
| | RPC5 | 275 T>M[23] | RPC5: K260, L276 | 73.1 | 11 | IV |

### Supplementary Table 3 | Cryo-EM data collection, refinement and validation statistics

| | Map A (EMDB-xxxx) Apo Pol III (PDB xxxx) | Map B (EMDB-xxxx) EC-1 Pol III (PDB xxxx) | Map C (EMDB-xxxx) | Map D (EMDB-xxxx) EC-3 Pol III (PDB xxxx) |
|---|---|---|---|---|
| **Data collection and processing** | | | | |
| Magnification | 105,000 | 130,000 | 130,000 | 130,000 |
| Voltage (kV) | 300 | 300 | 300 | 300 |
| Electron exposure (e–/Å$^2$) | 38.11 | 40.35 | 40.35 | 40.35 |
| Defocus range (μm) | 0.75-2.25 | 0.75-2.00 | 0.75-2.00 | 0.75-2.00 |
| Pixel size (Å) | 0.822 | 1.05 | 1.05 | 1.05 |
| Symmetry imposed | C1 | C1 | C1 | C1 |
| Initial particle images (no.) | 304,683 | 367,717 | 367,717 | 367,717 |
| Final particle images (no.) | 111,289 | 166,071 | 70,492 | 30,525 |
| Map resolution (Å) | 3.3 | 2.8 | 3.1 | 3.1 |
| FSC threshold | 0.143 | 0.143 | 0.143 | 0.143 |
| Map resolution range (Å) | 3.0 – 8.3 | 2.5 – 7.8 | 2.8 – 8.3 | 2.9 – 6.9 |
| **Refinement** | | | | |
| Initial model used (PDB code) | 3AYH,5AFQ, 5FJ8,5FJ9, 5FLM,5IY6, 6CND,6EU0, 6EU1,6EU2 | 3AYH,5AFQ, 5FJ8,5FJ9, 5FLM,5IY6, 6CND,6EU0, 6EU1,6EU2 | | 3AYH,5AFQ, 5FJ8,5FJ9, 5FLM,5IY6, 6CND,6EU0, 6EU1,6EU2 |
| Model resolution (Å) | 3.4 | 3.0 | | 3.2 |
| Map sharpening *B* factor (Å$^2$) | -80.1 | -55.2 | | -49.6 |
| **Model composition** | | | | |
| Non-hydrogen atoms | 40322 | 42712 | | |
| Protein / Nucleotide residues | 5072 / - | 5258 / 44 | | 5247 / 44 |
| Ligands | 1x SF4, 7x Zn, 1x Mg | 1x SF4, 7x Zn, 1x Mg | | 1x SF4, 7x Zn, 1x Mg |
| ***B* factors (Å$^2$)** | | | | |
| Protein | 105.65 | 84.75 | | 126.86 |
| Ligand | 105.80 | 93.50 | | 132.15 |
| Nucleotide | | 155.90 | | 180.37 |
| **R.m.s. deviations** | | | | |
| Bond lengths (Å) | 0.010 | 0.008 | | 0.008 |
| Bond angles (°) | 1.271 | 1.315 | | 1.208 |
| **Validation** | | | | |
| MolProbity score | 1.54 | 1.28 | | 1.43 |
| Clashscore | 3.91 | 2.20 | | 3.46 |
| Poor rotamers (%) | 0.25 | 0.43 | | 0.32 |
| **Ramachandran plot** | | | | |
| Favored (%) | 94.71 | 95.90 | | 95.71 |
| Allowed (%) | 5.23 | 3.99 | | 4.23 |
| Disallowed (%) | 0.06 | 0.12 | | 0.06 |

**Supplementary Table 3 | Cryo-EM data collection, refinement and validation statistics (continued)**

| | Map E (EMDB-xxxx) | Map F (EMDB-xxxx) | Map G* (EMDB-xxxx) | Map H* (EMDB-xxxx) EC-2 Pol III (PDB xxxx) |
|---|---|---|---|---|
| **Data collection and processing** | | | | |
| Magnification | 130,000 | 130,000 | 105,000/ 130,000 | 105,000/ 130,000 |
| Voltage (kV) | 300 | 300 | 300 | 300 |
| Electron exposure (e–/Å$^2$) | 40.35 | 40.35 | 38.11/40.35 | 38.11/40.35 |
| Defocus range (μm) | 0.75-2.00 | 0.75-2.00 | 0.75-2.25/ 0.75-2.00 | 0.75-2.25/ 0.75-2.00 |
| Pixel size (Å) | 1.05 | 1.05 | 1.05 | 1.05 |
| Symmetry imposed | C1 | C1 | C1 | C1 |
| Initial particle images (no.) | 367,717 | 367,717 | 277,360 | 277,360 |
| Final particle images (no.) | 100,152 | 21,861 | 277,360 | 31,737 |
| Map resolution (Å) | 2.9 | 3.3 | 2.9 | 3.3 |
| FSC threshold | 0.143 | 0.143 | 0.143 | 0.143 |
| Map resolution range (Å) | 2.6 – 7.6 | 3.0 – 8.9 | 2.8 – 7.1 | 3.2 – 10.0 |
| **Refinement** | | | | |
| Initial model used (PDB code) | | | | 3AYH,5AFQ, 5FJ8,5FJ9, 5FLM,5IY6, 6CND,6EU0, 6EU1,6EU2 |
| Model resolution (Å) | | | | 3.4 |
| Map sharpening *B* factor (Å$^2$) | | | | -43.6 |
| **Model composition** | | | | |
| Non-hydrogen atoms | | | | 42528 |
| Protein / Nucleotide residues | | | | 5233 / 44 |
| Ligands | | | | 1x SF4, 7x Zn, 1x Mg |
| ***B* factors (Å$^2$)** | | | | |
| Protein | | | | 117.34 |
| Ligand | | | | 143.32 |
| Nucleotide | | | | 222.81 |
| **R.m.s. deviations** | | | | |
| Bond lengths (Å) | | | | 0.006 |
| Bond angles (°) | | | | 1.155 |
| **Validation** | | | | |
| MolProbity score | | | | 1.44 |
| Clashscore | | | | 3.48 |
| Poor rotamers (%) | | | | 0.28 |
| **Ramachandran plot** | | | | |
| Favored (%) | | | | 95.64 |
| Allowed (%) | | | | 4.30 |
| Disallowed (%) | | | | 0.06 |

*Map G and map H derived from a combined data set of apo and elongating Pol III by merging final particle images of map A and map B. Data collection and processing values for both data sets are given.

**Supplementary Information references**

1. Davis, R. B., Likhite, N., Jackson, C. A., Liu, T. & Yu, M. C. Robust repression of tRNA gene transcription during stress requires protein arginine methylation. *Life Sci. Alliance* **2,** e201800261 (2019).

2. Eddy, S. R. Accelerated Profile HMM Searches. *PLoS Comput. Biol.* **7,** e1002195 (2011).
